# A time-resolved multi-omics atlas of transcriptional regulation in response to high-altitude hypoxia across whole-body tissues

**DOI:** 10.1101/2023.10.25.563964

**Authors:** Ze Yan, Ji Yang, Wen-Tian Wei, Ming-Liang Zhou, Dong-Xin Mo, Xing Wan, Rui Ma, Mei-Ming Wu, Jia-Hui Huang, Ya-Jing Liu, Feng-Hua Lv, Meng-Hua Li

## Abstract

High-altitude hypoxia acclimatization requires whole-body physiological regulation in highland immigrants, but the underlying genetic mechanism has not been clarified. Here we used sheep as an animal model for plain-to-plateau transplantation. We generated multi-omics data including time-resolved bulk RNA-Seq, ATAC-Seq and single-cell RNA-Seq from multiple tissues as well as phenotypic data from 20 bio-indicators. We characterized transcriptional changes of all genes in each tissue, and examined multi-tissue temporal dynamics and transcriptional interactions among genes. In particular, we identified critical functional genes regulating the short response to hypoxia in each tissue (e.g., *PARG* in the cerebellum and *HMOX1* in the colon). We further identified TAD-constrained *cis*-regulatory elements, which suppressed the transcriptional activity of most genes under hypoxia. Phenotypic and transcriptional evidence indicated that antenatal hypoxia could improve hypoxia tolerance in offspring. Furthermore, we provided time-series expression data of candidate genes associated with human mountain sickness (e.g., *BMPR2*) and high-altitude adaptation (e.g., *HIF1A*). Our study provides valuable resources and insights for future hypoxia-related studies in mammals.

## Introduction

Hypoxia is a severe challenge to an organism’s homeostatic equilibrium, affecting the physiological and pathological processes of organisms on plateaus^1,2^. The genetic mechanisms underlying long-term hypoxia adaptation have revealed positively selected genes and non-coding variants associated with cardiovascular, respiratory and metabolic traits in highland human and other vertebrates^3–6^. However, in human and animals inhabiting lowlands, visible physiological adjustments in response to hypoxia (e.g., acute increase in ventilation) occur during short-term acclimatization after moving to a plateau^7^, whose mechanisms have not yet been elucidated. In the case of maladaptation, hypoxia can lead to high-altitude diseases, such as polycythemia, pulmonary hypertension and heart failure in immigrants from lowland areas and chronic mountain sickness in high-altitude inhabitants^8–10^. Hypoxia also induces serious sickness in livestock transported to high-altitude areas, such as pulmonary hypertension in sheep^11^ and brisket disease in cattle^12^.

Sheep (*Ovis aries*) are an excellent model for studying hypoxia adaptation and acclimatization since they have adapted to a variety of environments (e.g., lowland and plateau) around the world^13,14^. Compared with other large animals, such as non-human primates, dogs, pigs and horses, sheep are an applicable animal model for biomedical research in terms of cost-effectiveness and ethical isssues^15,16^. For example, sheep models have been widely used for studying human cardiopulmonary, respiratory, neurological, immunological and reproductive diseases^17–21^ and for foetal-neonatal development and pathologies^22^. Additionally, Dolly the sheep was the first animal to ever be successfully cloned from cultured somatic cells^23^, making it possible to use the somatic cell nuclear transfer (SCNT) technique for both biomedical and agricultural applications of sheep^24^.

In this study, we performed a plain-to-plateau transplantation experiment in sheep (Fig. 1a) and generated multiple layers of data (Fig. 1b), including whole-body transcriptomes, chromatin accessibility (i.e., ATAC-Seq) and single-cell RNA-Seq (scRNA-Seq) data, and blood physiological and biochemical phenotypes. We aimed to (1) identify time-series expression changes and regulatory elements in response to short-term hypoxia across tissues; (2) reveal multi-tissue expression patterns of genes implicated in high-altitude adaptation and diseases in human; and (3) test the inheritance of acclimatization to hypoxia between generations. This research promotes our understanding of the mechanisms conferring resilience to hypoxia and provides valuable resources for future studies of hypoxia-related diseases in human and livestock.

**Fig. 1.**
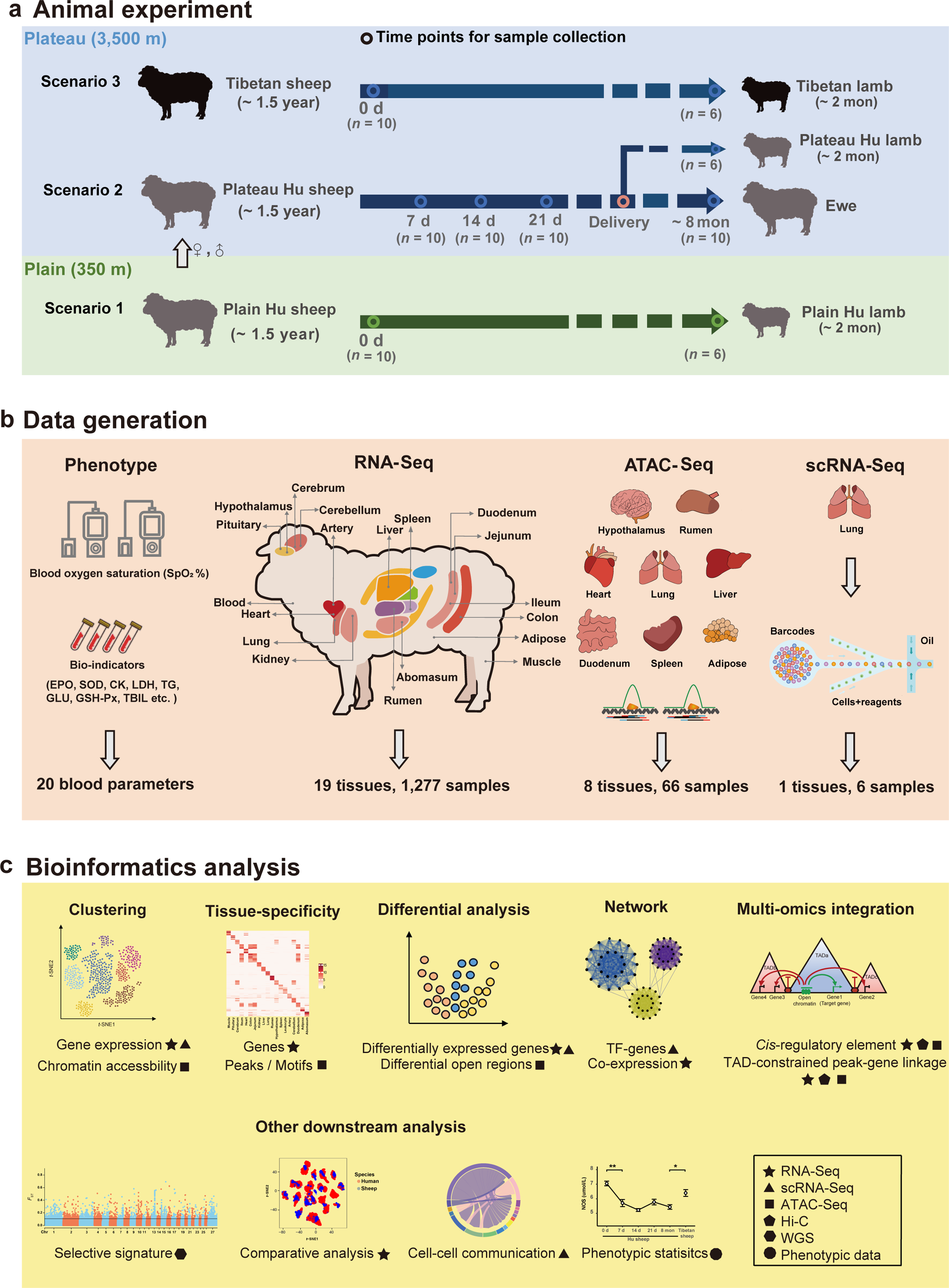
Schematic diagram of the study. a, Design of the animal transplantation experiment. Hu sheep (grey) and Tibetan sheep (black) originally inhabited plain and plateau environments, respectively. There were three scenarios examined in our experiment: plain Hu sheep raised in lowlands (scenario 1); plateau Hu sheep, namely plain Hu sheep transplanted to highlands (scenario 2) and acclimatized until four time points; and Tibetan sheep raised in highlands (scenario 3). In addition, offspring of the ewes in the above three scenarios were included. b, Sample collection and data generation. We collected 19 whole-body tissues and produced phenotypic, transcriptomic (bulk-RNA, single-cell RNA) and epigenetic (ATAC-Seq) data. c, Major bioinformatics and statistical analysis involved in the study.

## Results

### Plain-to-plateau transplantation experiment

Hu sheep and Tibetan sheep, two representative Chinese native breeds that originally inhabited plains (Zhejiang Province, China) and the Qinghai-Tibet Plateau (QTP), respectively, were included in the experiment. Tibetan sheep initially spread to the QTP from northern China along with the colonization of nomads ∼3,100 years ago and have become well adapted to the high-altitude environment^25^. There were three different scenarios in our experiment (Fig. 1a): (*i*) scenario 1, plain Hu sheep (i.e., ewes of Hu sheep the plain, *n* = 10) raised in Neijiang (∼ 350 m.a.s.l, the southeast of Sichuan Province, “0 d” hereafter); (*ii*) scenario 2, plateau Hu sheep (*n* = 43: ewes, *n* = 40; rams, *n* = 3) that were transplanted from the plain to Aba Autonomous Prefecture (∼3,500 m.a.s.l, eastern edge of the QTP in Sichuan Province) and acclimatized for four different time periods after transplantation (i.e., 7 days, 14 days, 21 days and ∼8 months, “7 d, 14 d, 21 d, 8 mon” hereafter); and (*iii*) scenario 3, Tibetan sheep (i.e., ewes of Tibetan sheep on the plateau; *n* = 10) raised in Aba. In addition, offspring (*n* = 18) of the ewes under the above three scenarios (six lambs for each scenario) were included in our experiment.

### Data summary

To comprehensively study transcriptomic, epigenomic and phenotypic changes in sheep acclimatization from the plain to the plateau, we collected multiple tissues from experimental animals to perform high-throughput sequencing (Fig. 1b). We produced 1,277 RNA-Seq datasets from 19 major tissues (Supplementary Tables 1 and 2) and then uniformly processed the RNA-Seq datasets to yield ∼ 23 billion uniquely mapped paired-end reads, with a mean mapping rate of 84.3% (61.05-91.37%) (Supplementary Table 3). We also generated 66 chromatin accessibility datasets for eight tissues (i.e., heart, artery, lung, liver, hypothalamus, rumen, duodenum and adipose) by ATAC-Seq, and we produced six high-resolution scRNA-Seq datasets for lung tissue (i.e., for plain Hu sheep, plateau Hu sheep at four acclimatization time points and Tibetan sheep) (Supplementary Tables 1 and 2). After raw data processing, we obtained ∼ 4.5 billion informative reads with an average unique mapping rate of 98.63% (84.57% - 99.27%) for ATAC-Seq (Supplementary Table 4) and ∼ 4.0 billion reads with a confident mapping rate of 79.71% (52.70% - 95.40%) for scRNA-Seq (Supplementary Table 5).

To evaluate physiological acclimatization under high-altitude hypoxia, we collected phenotypic data for 20 blood parameters, including blood oxygen saturation (SpO_2_) and 19 bio-indicators (e.g., erythropoietin, nitric oxide and cardiac enzymes) (Supplementary Tables 6 and 7). We also included 37 whole-genome sequences^26^ and high-throughput chromosome conformation capture (Hi-C) data from a sheep that were previously published^27^ for the integrated analysis (Fig. 1c and Supplementary Tables 8 and 9).

### Phenotypic and transcriptional characteristics

Previous evidence from several vertebrate taxa suggested that physiological adjustments have played a significant role in high-altitude hypoxia tolerance and could well represent environment-induced physiological changes^3,28^. We examined 10 ewes of Hu sheep before (i.e., 0 d) and after their transplantation to the QTP at four time points (i.e., 7 d, 14 d, 21 d and ∼8 mon) to track the changes in SpO_2_ and other blood indicators. We observed that the mean value of SpO_2_ decreased sharply (0 d vs. 7 d, Wilcoxon rank sum test, *P* = 7.50 ×10^−13^) at 7 d but increased constantly at 14 d and later with acclimatization (Fig. 2a). However, compared to Tibetan sheep, Hu sheep still exhibited significantly lower levels of SpO_2_ (Hu sheep: 85.92; Tibetan sheep: 87.71; Wilcoxon rank sum test, *P* = 1.40 ×10^−9^) even after 8 months (Fig. 2a), probably reflecting potentially different mechanisms underlying phenotypic plasticity and genetic adaptation to high-altitude hypoxia in mammals^3^. In addition to SpO_2_, we observed distinct patterns of the other bioindicators in the transplanted Hu sheep (Extended Data Fig. 1). For example, nitric oxide synthetase (NOS), which restricts the synthesis of the vasodilator NO, was decreased after transplantation and showed significantly lower average values than in Tibetan sheep (Hu sheep: 5.43 µmol/L, Tibetan sheep: 6.40 µmol/L; Wilcoxon rank sum test, *P* =0.013) (Fig. 2b), verifying that down-regulated NO synthesis contributes to hypoxic pulmonary vasoconstriction^28–30^. In general, the levels of triglycerides (TG) and glucose (GLU), which are associated with energy metabolism, increased over time (Extended Data Fig. 1), suggesting enhanced energy production in response to hypoxia. The change in total bilirubin (T-BIL), an indicator that is positively correlated with liver damage, followed a bell curve (Extended Data Fig. 1), implying that liver impairment was gradually relieved during acclimatization. Cardiac enzymes (CK) showed non-significant changes among the four time points examined during acclimatization (*P* > 0.05) but presented significantly lower averages than in Tibetan sheep (Hu sheep: 58.96 U/L; Tibetan sheep: 146.7 U/L; Wilcoxon rank sum test, *P* = 0.0087) (Fig. 2c), indicating nonactivation during short-term hypoxia.

**Fig. 2.**
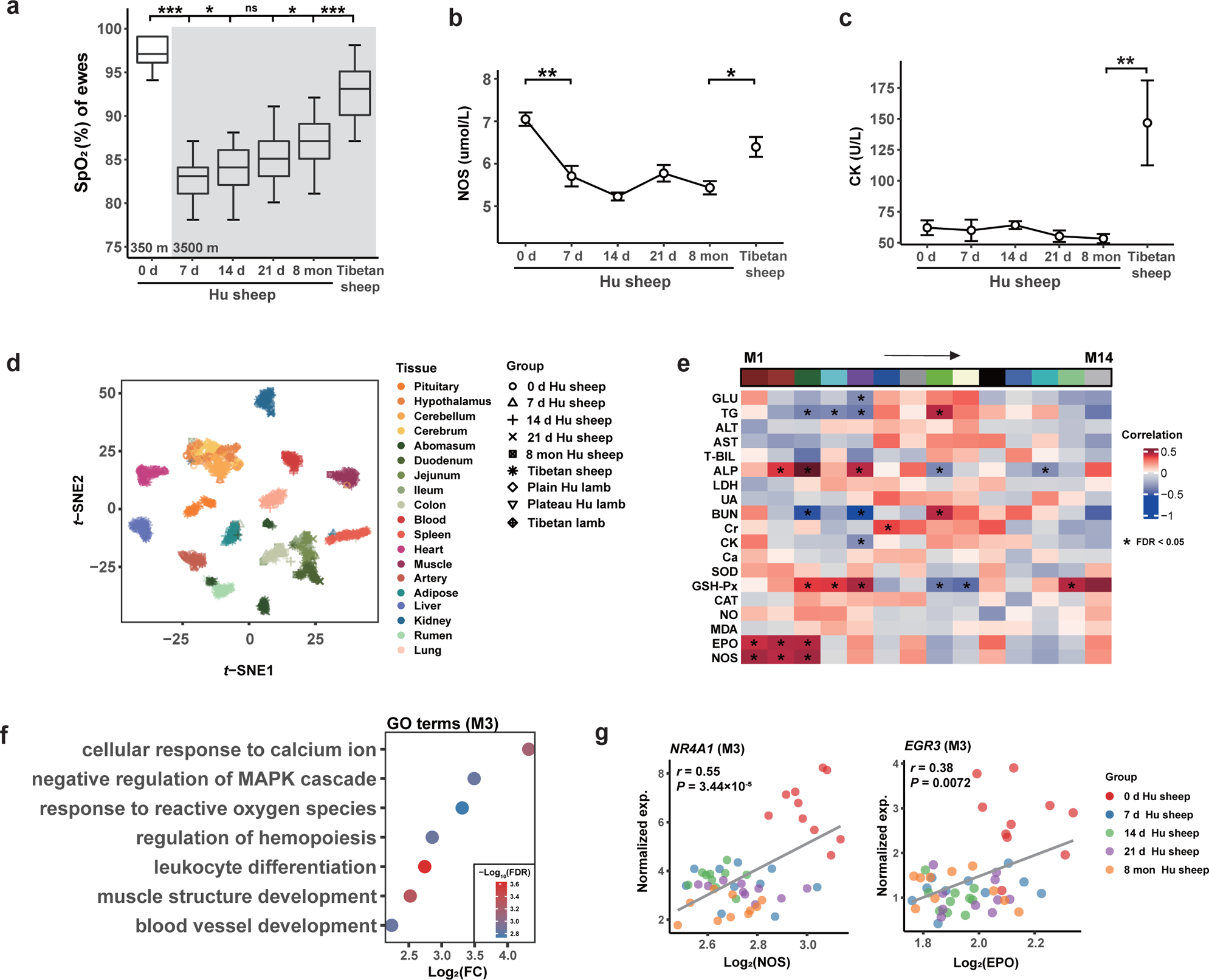
Characteristics of phenotype and gene expression. a, Changes in blood oxygen saturation (SpO_2_) with acclimatization time. b, Changes in the nitric oxide synthetase (NOS) value over time. c, Similar to b, changes in cardiac enzyme (CK) values over time. *P* values from the Wilcoxon rank sum test, * *P* < 0.05, ** *P* < 0.001, *** *P* < 0.0001, and “ns” means not significant. d, *t*-distributed stochastic neighbour embedding (*t*-SNE) clustering of 1,277 RNA-Seq samples based on normalized expression (transcripts per million, TPM). e, Association between gene modules and bio-indicators in blood using weighted gene co-expression network analysis (WGCNA). The rows represent the 14 gene modules (i.e., M1-M14), and the columns show 19 bio-indicators. Multiple testing was corrected using the Benjamini-Hochberg method. * FDR < 0.05. f, Gene Ontology (GO) enrichment analysis for gene module 3 (M3). g, Gene examples (i.e., *NR4A1* and *EGR3*) in M3. Scatter plots show the Pearson’s correlation between the expression levels of genes and the values of bioindicators over time.

To characterize the global expression patterns of whole-body tissues, we first applied *t*-distributed stochastic neighbour embedding (*t*-SNE) analysis based on the gene expression profiles of samples. The resultant sample clustering recapitulated the different tissues accurately (Fig. 2d), consistent with the hierarchical clustering of expression profiles (Extended Data Fig. 2a-d). The functions of the genes with tissue-specific expression reflected known tissue biology (Supplementary Table 10).

Next, we explored the effect of hypoxia on gene expression within tissues over time. We only observed a typical pattern of gene expression in the abomasum, which showed an obvious separation of samples before and after 14 d (Extended Data Fig. 3). In particular, *GKN2,* an abomasum-specific gene that is associated with oxidative stress-induced gastric cancer cell apoptosis^31,32^, showed increased expression with time (Extended Data Fig. 4).

Furthermore, we investigated associations between blood gene expression and bio-indicators using weighted correlation network analysis (WGCNA). Among 13,707 genes remaining after filtration, we determined 14 gene modules (in M1-M14) labelled with different colours (Extended Data Fig. 5a, b), and 10 gene modules were significantly correlated (FDR < 0.05) with the bio-indicators (Fig. 2e, Extended Data Fig. 5c and Supplementary Table 11). In particular, we observed that M3 was significantly positively associated with erythropoietin (EPO), nitric oxide synthetase (NOS), glutathione peroxidase (GSH-Px) and alkaline phosphatase (ALP). The results of Gene Ontology (GO) enrichment among the genes in M3 were congruent with their associated bio-indicators (Fig. 2f). For example, EPO is involved in the regulation of hemopoiesis (e.g., *EGR3*, *HOX5* and *FOS*), NOS is associated with blood vessel development (e.g., *NR4A1*, *JUN* and *RHOB*), and GSH-Px is related to the response to reactive oxygen species (e.g., *NR4A3*, *PLK3* and *TNFAIP3*). Protein-protein interaction analysis also showed that genes (e.g., *JUN*, *FOS* and *NR4A1*) related to the above GO terms were at the centre of the regulatory network within the genes in M3 (Extended Data Fig. 5d). Furthermore, we explored the changes between gene expression and bio-indicators over time (Extended Data Fig. 6). The overall expression levels of *NR4A1* and *EGR3* (from gene module M3 in Fig. 2e) were significantly and positively correlated with NOS (Pearson’s *r* = 0.55, *P* = 3.44 × 10^−5^) (Fig. 2g) and with EPO (Pearson’s *r* = 0.38, *P* = 0.0072) (Fig. 2g), respectively. *NR4A1* showed higher expression and higher NOS values at 0 d (i.e., normoxia) (Extended Data Fig. 5e). A similar expression trend was also observed for *EGR3* (Extended Data Fig. 5f).

### Temporal transcriptome dynamics and multi-tissue interactions

To explore the transcriptional changes during hypoxia acclimatization, we identified differentially expressed genes (DEGs) between five adjacent time points across tissues. Overall, we observed large variations in transcriptional regulation between and within tissues in terms of the number of DEGs, particularly between 0 d and 7 d (Fig. 3a). We identified the most active tissues (i.e., tissues with the top 3 counts of DEGs) in each comparison. Certain tissues (e.g., kidney, colon, adipose and cerebellum) showed activation in the comparisons of multiple or, particularly, adjacent time points (Fig. 3a). For example, kidney showed activation in all four comparisons of adjacent time points, indicating that kidney functions concerning ATP production and stress hormone secretion were important for the hypoxia response^33,34^. Colon showed activation in both the “7 d vs. 14 d” and “14 d vs. 21 d” comparisons, implying that hypoxia strongly affects intestinal homeostasis^35^ during acclimatization. We also found that the cerebellum only showed activation in “0 d vs. 7 d”, which suggested that hypoxia severely affects the cerebellum first, before the other examined tissues. These observations demonstrated that certain organs or tissues, such as kidney, colon, adipose and cerebellum, that were actively involved in rapid hypoxia acclimatization had main functions (e.g., energy metabolism, endocrine, and nervous system functions) that differed from those of tissues (e.g., heart and lung) involved in long-term hypoxia adaptation, such as cardiovascular and respiratory functions.

**Fig. 3.**
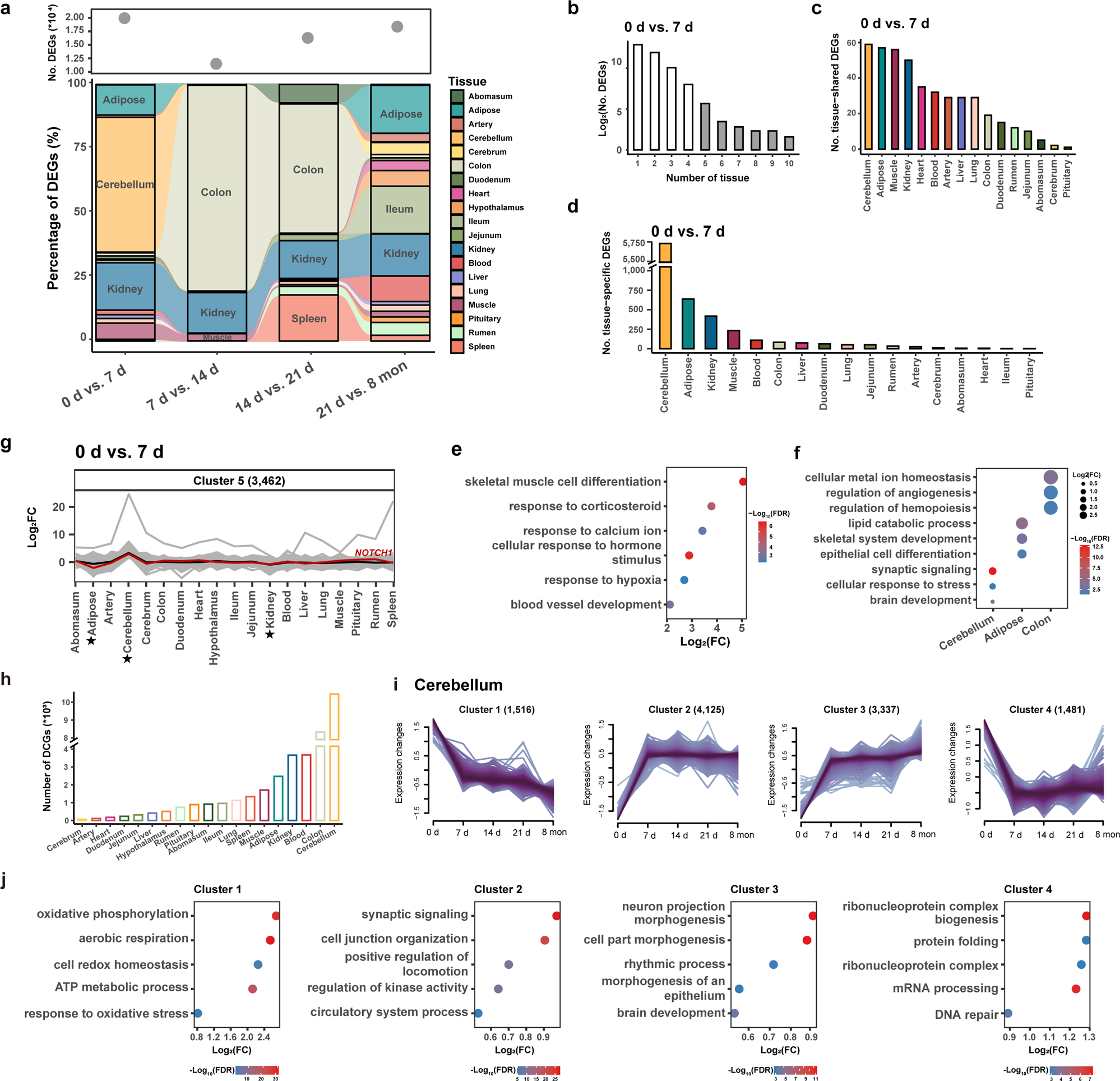
Transcriptome dynamics during hypoxia acclimatization. a, Numbers of differentially expressed genes (DEGs) (top) and percentages of DEGs (bottom) between adjacent time points comparisons across tissues. b, Distribution of DEGs across numerous tissues in the “0 d vs. 7 d” comparison. c-d, Numbers of tissue-shared and tissue-specific (d) DEGs across tissues in the “0 d vs. 7 d” comparison. e-f, GO term enrichments for tissue-shared (e) and tissue-specific (f) DEGs from the “0 d vs. 7 d” comparison. g, Multi-tissue interactions in the “0 d vs. 7 d” comparison. The average log_2_FC value for cluster 5 is denoted with a black line. Active tissues (i.e., cerebellum, kidney and colon) are marked with asterisks, and the interaction of *NOTCH* between tissues is highlighted with a red line. h, Numbers of dynamically changed genes (DCGs) across tissues. i, Fuzzy *c*-means clustering identified gene expression patterns of DCGs in the cerebellum. j, GO terms for the four clusters identified in i.

Next, we examined the distribution of DEGs across tissues for each comparison. Most of the DEGs were assigned to particular tissues, while a small number of DEGs showed a ubiquitous distribution (Fig. 3b and Extended Data Fig. 7a-c). We then identified tissue-shared (i.e., in at least five tissues) and tissue-specific DEGs (i.e., in only one tissue) for each comparison (Fig 3c, d and Supplementary Tables 12 and 13). In the comparison of 0 d vs. 7 d, the functional enrichment of tissue-shared DEGs showed the involvement of the genes in multiple biological processes (Supplementary Table 14), such as skeletal muscle cell differentiation (e.g., *BTG2*, *ATF3* and *NR4A1*), response to hypoxia (e.g., *NR4A2*, *EGR1* and *CPEB2*) and the response to corticosteroids (e.g., *NR4A3*, *IGF1R* and *FOS*) (Fig. 3e). However, tissue-shared DEGs from the other three comparisons mostly participated in energy metabolism, such as mitochondrial organization (e.g., *NDUFAF8*, *ROMO1* and *UQCC2*) and ATP metabolic processes (e.g., *ND1*, *COX2* and *ATP5PF*) (Supplementary Table 14). These results implied that the initial hypoxic stimulus (e.g., the first 7 d after transplantation to QTP) resulted in collective responses of multiple life systems^4^. Additionally, the functions of tissue-specific DEGs reflected the respective tissue biology and hypoxia response (Supplementary Table 14). For instance, in the comparison of “0 d vs. 7 d”, cerebellum-specific DEGs were associated with synaptic signalling (e.g., *NTNG1*, *TNF* and *GABBR2*) and regulation of the cellular response to stress (*HIF1A*, *ATF4* and *PARG*), while colon-specific DEGs were involved in cellular metal ion homeostasis (e.g., *HMOX1*, *TRPM8* and *CXCR5*) and the regulation of angiogenesis (e.g., *ISL1*, *THBS4* and *ADGRB2*) (Fig. 3f). Taken together, the findings indicated that hypoxia acclimatization could have activated both hypoxia response processes in multiple tissues and the functions of specific tissues, which were collectively regulated by polygenic (i.e., tissue-specific DEGs) and pleiotropic (i.e., tissue-shared DEGs) genes^36^.

Previous evidence suggests that the maintenance of systemic homeostasis and responses to environmental challenges typically requires transcriptional interactions among multiple organs and tissues ^37,38^. To identify transcriptional interactions underlying hypoxia acclimatization, we used the *k*-means method to analyse multi-tissue interactions based on log_2_FC (log_2_-transformed fold change) values in the above differential expression analysis and obtained 5-7 clusters in different comparisons (Extended Data Fig. 8a-d). Overall, in the comparison of “0 d and 7 d”, gene cluster 5 (e.g., *NOTCH1*) showed the greatest increase in expression in the cerebellum but decreased expression in kidney and adipose at 7 d compared to 0 d (Fig. 3g and Extended Data Fig. 8e). Notably, we observed possible interactions among the most active tissues, such as cerebellum, kidney and adipose in the “0 d vs. 7 d” comparison (Fig. 3a) and colon in the “7 d vs. 14 d” and “14 d vs. 21 d” comparisons (Extended Data Fig. 8b, c). These findings implied the action of potential transcriptional networks among particular tissues at different time points during acclimatization.

### Time-series expression changes

To explore the expression patterns within tissues across time points, we conducted a time-series differential expression analysis to identify dynamically changed genes (DCGs) (i.e., genes with significant expression changes throughout the acclimatization process). Since genes with similar expression patterns could be involved in the same biological process^39,40^, we further classified DCGs into different gene clusters based on their expression patterns with the *c*-means method^41^ (Supplementary Table 15). The numbers of our DCGs in different tissues ranged from 68 (cerebrum) to 10,459 (cerebellum) (Fig. 3h) and were categorized into 2-6 clusters across tissues (Extended Data Fig. 9). In most tissues, the changes in the expression of DCGs over time reflected similar patterns of the temporal transcriptional changes described above (Fig. 3a and Extended Data Fig. 9). For example, DCGs in the cerebellum were categorized into four clusters (Fig. 3i), and the overall expression patterns of these clusters varied greatly at 7 d. This observation was consistent with the large transcriptional changes in the cerebellum in the “0 d vs. 7 d” comparison (Fig. 3a). Furthermore, the DCGs in each cluster from the cerebellum exhibited distinct biological functions (Fig. 3j). Specifically, the functions of the DCGs in cluster 1 were associated with energy metabolism (e.g., aerobic respiration and ATP metabolic process), while in cluster 3, the gene functions were related to brain biology (e.g., neuron projection morphogenesis and brain development) (Fig. 3j). These results revealed adjustments in energy metabolism and the biological function of the cerebellum in response to hypoxic challenge.

### Hypoxia-adaptive genes in adaptation and acclimatization

The evolution of gene expression and regulation is a major source of phenotypic diversity ^42–44^. To explore the roles of hypoxia-adaptive genes in long-term adaptation and short-term acclimatization, we integrated gene expression data with genome sequencing data to perform a joint analysis. We first calculated pairwise *F*_ST_ values between 37 genomes of 7 sheep breeds originating from the plain (*n* = 20) and plateau (*n* = 17) regions and chose the top 5% of the *F*_ST_ distribution as candidate selected regions (Extended Data Fig. 10a and Supplementary Table 16). The functional annotation of putatively selected genes (i.e., *F*_ST_ genes) from the candidate regions revealed their high relevance to high-altitude adaptation (Extended Data Fig. 10b and Supplementary Table 17). We detected DEGs between plain Hu sheep and Tibetan sheep in each of the tissues (Supplementary Table 18). For each tissue, we intersected *F*_ST_ genes with inter-breed DEGs and DCGs separately. We examined the distribution of two categories of intersected genes (i.e., *F*_ST_ genes in DEGs and *F*_ST_ genes in DCGs) across tissues (Fig. 4a, b and Supplementary Table 19). We identified 52 multi-tissue *F*_ST_ genes (i.e., *F*_ST_ genes in at least five tissues) in DEGs and 179 multi-tissue *F*_ST_ genes in DCGs, including 13 common genes (i.e., *APOLD1*, *NDUFB9*, *ERBB4*, *NFKBIZ*, *NR4A3*, *RPS8*, *CIAO2A*, *AHCYL2*, *ESRRG*, *KIAA0930*, *RASGEF1B*, *MRPS25* and *TNFRSF21*) (Fig. 4c and Supplementary Table 20). The results indicated that these 13 *F*_ST_ genes could have played an important role in hypoxia adaptation and acclimatization by regulating the expression of multiple tissues. Furthermore, we examined changes in the expression levels of these genes across sheep tissues (Fig. 4d and Extended Data Fig. 11) and investigated their functions in the human GWAS atlas database (Supplementary Table 21). We found that the human trait/disorder associations (e.g., hypoxia-related traits) of these genes were largely consistent with dynamic expression changes in analogous sheep tissues. For instance, *APOLD1*, which was significantly (*P* < 0.05) associated with cardiovascular (e.g., high blood pressure), respiratory (e.g., asthma) and haematological (e.g., haemoglobin) traits (Fig. 4e and Supplementary Table 21), showed significant expression changes in the heart, lung and kidney (Fig. 4d). Likewise, *NR4A3* was significantly (*P* < 0.05) associated with metabolic (e.g., fat-free mass), nervous/neurological (e.g., neuroticism and insomnia) and cardiovascular (e.g., resting heart rate) traits (Fig. 4e and Supplementary Table 21) and was dynamically expressed in adipose, cerebellum, heart and artery (Fig. 4d). These results suggested that the 13 identified tissue-shared hypoxia-adaptive genes could regulate hypoxia-related traits by controlling expression in different tissues in both genetic adaptation and short-term acclimatization.

**Fig. 4.**
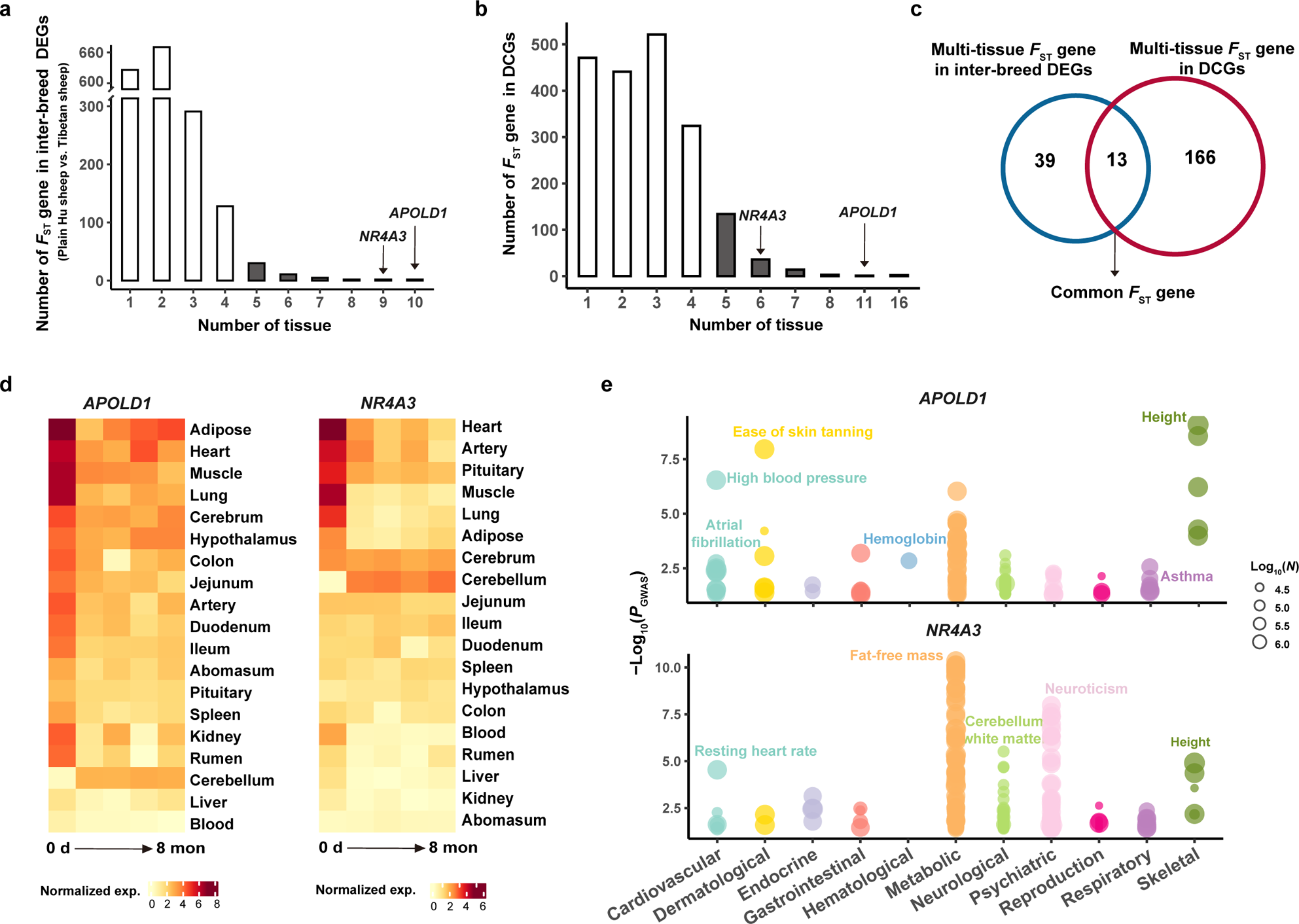
Hypoxia-adaptive genes in adaptation and acclimatization. a, Distribution of *F*_ST_ genes among differentially expressed genes (DEGs) between breeds (i.e., plain Hu sheep vs. Tibetan sheep) across tissues. b, Distribution of *F*_ST_ genes in DCGs across tissues. c, Venn diagram showing the intersection of multi-tissue *F*_ST_ genes in inter-breed DEGs with those in DCGs. d-e, Examples of common multi-tissue *F*_ST_ genes. d, Expression changes in *APOLD1* (left) and *NR4A3* (right) over time across tissues. e, Phenome-wide association analysis (Phe-WAS) for *APOLD1* (top) and *NR4A3* (bottom). *N* is the sample size of GWAS.

### Characterization of chromatin accessibility across tissues under hypoxia

To identify regulatory elements related to dynamic expression, we applied ATAC-Seq to detect genomic chromatin accessibility across eight important tissues (i.e., hypothalamus, rumen, heart, lung, liver, duodenum, spleen and adipose) under the three scenarios described above. We obtained a total of 1,662,152 statistically significant peaks (*P* < 0.01) (Extended Data Fig. 12a), and the open chromatin regions in the eight tissues were highly enriched at transcription start sites (TSS) (Fig. 5a). The distribution of peaks varied among tissues (Extended Data Fig. 12b), but in general, the highest proportion of peaks were located in intergenic regions, followed by intronic region, while the lowest proportion were located in 5’ UTRs (Extended Data Fig. 12c). One exception was adipose, which showed the largest number of peaks in regions ≤ 1 kb from promoters. The ATAC-Seq signals showed strong correlations among biological replicates at the whole-genome level (Extended Data Fig. 12d) and were clustered by different tissues instead of breeds and time points (Fig. 5b). Moreover, we observed a strong positive correlation (Pearson’s *r* = 0.66, *P* = 0.076) between the numbers of protein-coding genes (PCGs) and ATAC-Seq peaks across tissues (Fig. 5c), which demonstrated that open chromatin regions positively regulate transcriptional activity.

**Fig. 5.**
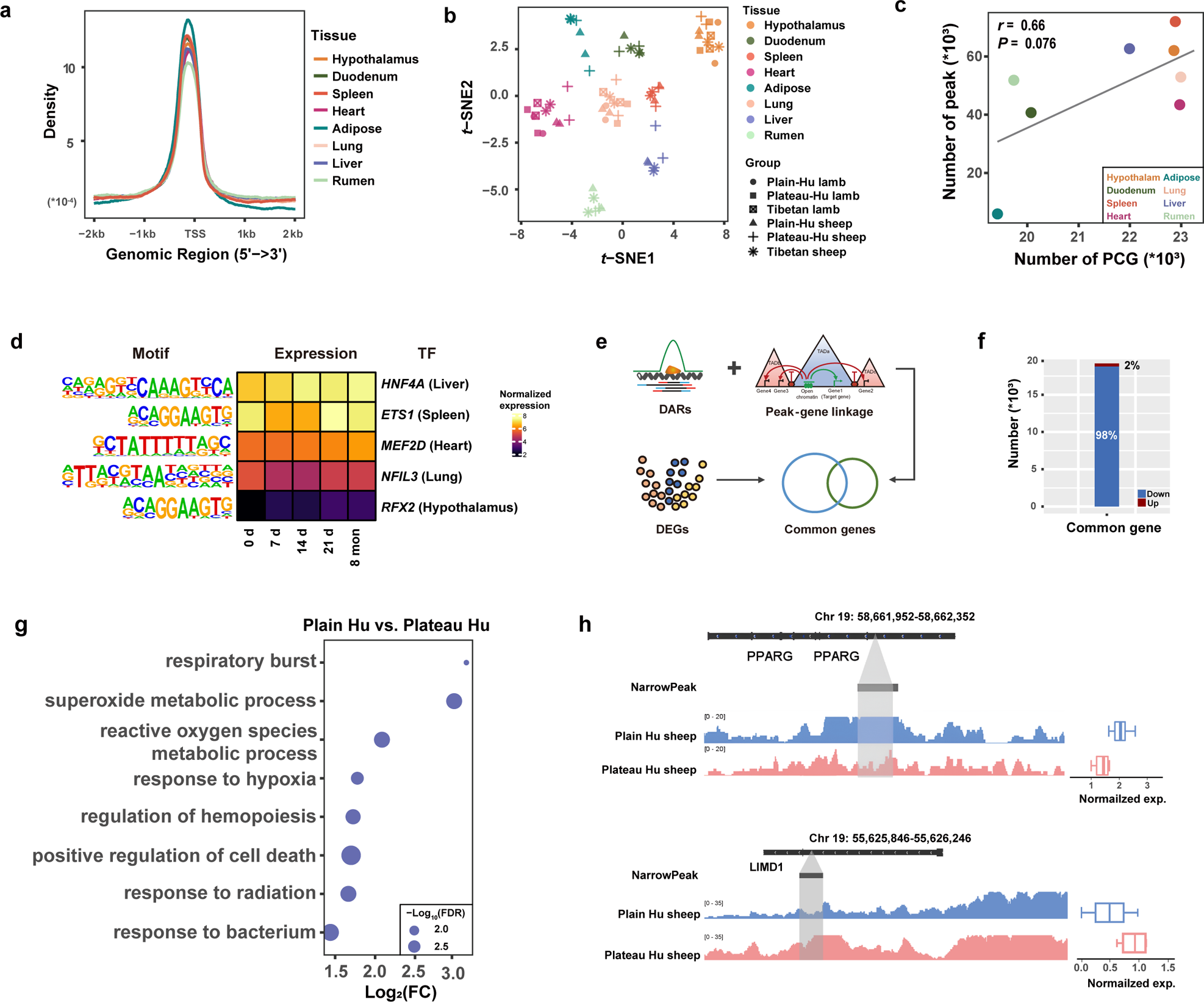
Chromatin accessibility reveals the regulatory landscape of hypoxia acclimatization. a, Average peak density of each tissue at positions relative to transcription start sites (TSSs). b, *t*-SNE clustering of 66 samples based on peak signal density. c, Pearson’s correlations between the numbers of expressed genes and detected peaks across tissues. PCGs: protein coding genes. d, Tissue-specific transcription factors (TFs) in differentially accessible regions (DARs) of the respective tissues and their gene expression over time. e, Identification of common genes. f, Numbers of up-regulated and down-regulated peak-gene pairs. g, Common genes of liver from plain Hu sheep vs. plateau Hu sheep comparison are enriched in biological processes related to the hypoxia response. h, Examples of common up-regulated and down-regulated genes. Peak density and expression level of the down-regulated common gene *PPARG* (top) in the liver and the up-regulated common gene *LIMD1* (bottom) in the hypothalamus from plain Hu sheep and plateau Hu sheep comparison.

As hotspots for transcription factor (TF) activities, open chromatin landscapes have unique effects in driving the biological functions of tissues^45,46^. We further characterized tissue-specific peaks and identified corresponding tissue-specific TFs (Extended Data Fig. 13a and Supplementary Table 22). Tissue-specific peaks were largely located in distal intergenic regions (Extended Data Fig. 13b), and TF binding motifs (TFBMs) were significantly (*P* < 0.05) enriched in the specific peaks of various tissues, such as the TFBMs of *MEF2D* in heart, *HNF4A* in liver and *ETS1* in spleen (Extended Data Fig. 13c,d). We also implemented differential expression analysis between pairwise comparisons of plain Hu sheep, plateau Hu sheep after transplantation to the QTP for 8 months and Tibetan sheep for each tissue and determined TFBMs in differentially accessible regions (DARs). We found that some tissue-specific TFBMs were significantly (*P* < 0.05) enriched in DARs of corresponding tissues (Fig. 5d and Supplementary Table 23). For example, the heart-specific TFBM of *MEF2D* was found in DARs of heart tissue. The expression of *MEF2D* gradually increased with time (Fig. 5d), suggesting the continuous activation of *MEF2D* target genes in heart, which is in line with the role of *MEF2D* in the regulation of cardiac muscle^47^.

### Regulation of gene expression by TAD-constrained *cis*-regulatory elements

To leverage the chromatin accessibility information captured by integrating gene expression, we used a correlation-based method to predict *cis*-regulatory elements (CREs) and their target genes within the same topologically associated domains (TADs), enabling the capture of all CREs (e.g., promoters and enhancers). A total of 2,875,658 independent peak-gene assignments were derived from all the TADs, and after filtration, 460,421 high-quality peak-gene pairs were retained for the following analysis. Subsequently, we examined the functions of target genes for the tissue-specific and conserved peaks. The target genes reflected tissue specificity and tissue biological functions well (Supplementary Table 24). For example, lung target genes were significantly (FDR < 0.05) associated with lung development (e.g., *FOXF1*, *NKX2-1* and *LIF*) and epithelial cell differentiation (e.g., *HOXA7*, *TMOD1* and *SOX17*) (Extended Data Fig. 13e). This observation indicated that CREs frequently interact within TADs to regulate gene expression.

We further explored the role of CREs in the regulation of gene expression during hypoxic acclimatization. We first performed differential expression analysis between pairwise comparisons of plain Hu sheep, plateau Hu sheep after transplantation to the QTP for 8 months and Tibetan sheep based on the RNA-Seq data. For each comparison, we annotated target genes linked to the DARs and detected the common genes showing both up- or down-regulated expression and changes in chromatin accessibility (Fig. 5e and Supplementary Table 25). We identified a total of 19,151 common peak-gene pairs between groups (i.e., plain Hu sheep vs. plateau Hu sheep, plain Hu sheep vs. Tibetan sheep and plateau Hu sheep vs. Tibetan sheep) across tissues, including 364 up-regulated and 18,787 down-regulated genes (Fig. 5f). We found that the common genes identified from the comparison of plain Hu sheep vs. plateau Hu sheep were related to hypoxia adaptation. For example, down-regulated common genes in plateau Hu sheep were significantly (FDR < 0.05) enriched in the response to hypoxia and regulation of hemopoiesis in the liver (Fig. 5g and Supplementary Table 26). In particular, *PPARG*, whose expression was down-regulated due to less accessible chromatin (Fig. 5h), is relevant to the regulation of cardiovascular circadian rhythms^48^. Additionally, *LIMD1*, whose functions are associated with the regulation of hippo signalling^49^ and the response to hypoxia^50^, was up-regulated in the hypothalamus in plateau Hu sheep due to open accessibility (Fig. 5h). Therefore, the expression of the aforementioned common genes was regulated (i.e., up- or down-regulated) by chromatin accessibility and further affected hypoxia acclimatization.

### Testing the heritability of acclimatization to hypoxia

To test the heritability of acclimatization to hypoxia, we examined the values of SpO_2_, gene expression and chromatin accessibility in lambs and ewes of the three sheep groups (i.e., plain Hu, plateau Hu and Tibetan sheep). Strikingly, the plateau Hu lambs showed no significant difference in the mean value of SpO_2_ measures from Tibetan lambs (plateau Hu sheep: 85.92, Tibetan sheep: 87.7, Wilcoxon rank sum test, *P* = 0.16) (Fig. 6a). We performed differential expression analysis between pairwise comparisons of the three lamb groups and focused on the DEGs from the comparisons of plateau Hu lambs vs. plain Hu lambs and Tibetan lambs vs. plain Hu lambs (Fig. 6b). Based on comparison with plain Hu lambs, we found that the DEGs detected in plateau Hu lambs and Tibetan lambs were significantly (FDR < 0.05) enriched in many common GO terms, such as extracellular matrix organization in kidney, localization within membranes in cerebrum and the cellular response to angiotensin in artery (Fig. 6c and Supplementary Table 27). Some of these GO terms (e.g., extracellular matrix organization and localization within membrane) were directly activated by hypoxia^51^, suggesting that plateau Hu lambs and Tibetan lambs may share similar hypoxia-responsive biological processes. Additionally, we noticed that several hypoxia response processes were only identified in the comparison of plateau Hu lambs vs. plain Hu lambs, including the response to hypoxia in lung and response to decreased oxygen levels in artery and lung (Fig. 6d).

**Fig. 6.**
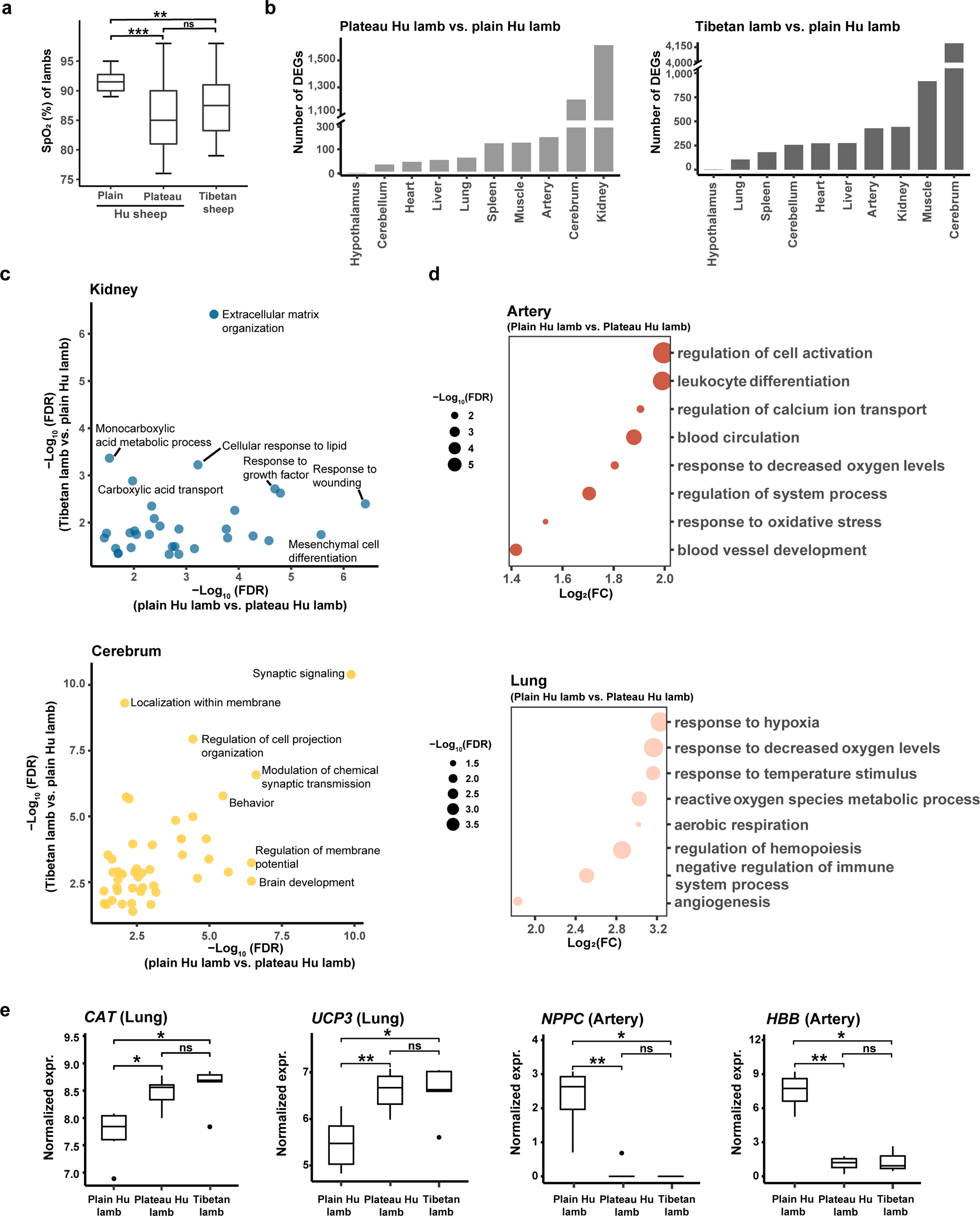
Hypoxia-exposed parents enhance the hypoxia tolerance of offspring. a, Changes in SpO_2_ in the three lamb groups. b, Numbers of DEGs from the plateau Hu lamb vs. plain Hu lamb (left) and Tibetan lamb vs. plain Hu lamb comparisons (right). c, Common GO terms enriched with the DEGs of plain Hu lambs vs. plateau Hu lambs and plain Hu lambs vs. Tibetan lambs in the kidney (top) and cerebrum (bottom). d, GO terms enriched only with the DEGs from the plain Hu lamb and plateau Hu lamb comparison in artery (top) and lung (bottom). e, Expression levels of key genes from hypoxia response-related GO terms. *P* values come from the Wilcoxon rank sum test, * *P* < 0.05, ** *P* < 0.001, *** *P <* 0.0001, and “ns” indicates not significant.

We further examined the expression profiles of the important functional genes *CAT* and *UCP3* in response to hypoxia in lung, *NPPC* in response to decreased oxygen levels and *HBB* in blood circulation in artery. The expression patterns of the genes were similar to the patterns of SpO_2_ alterations (Fig. 6e). Moreover, we identified common genes showing both up- or down-regulated expression and changes in chromatin accessibility between the lamb groups for each sampled tissue (i.e., lung, heart, hypothalamus) (Supplementary Table 25). For example, the expression of functional genes for high-altitude adaptation, such as *SIK1, OTOF, SOCS1* and *JUN* in heart and *CXCL8*^52–54^ in lung, showed significant (*P* < 0.05) downregulation in plateau Hu lambs compared to plain Hu lambs. Overall, the results indicated that plateau Hu lambs show developed adaptive characteristics at birth according to the three above measures, presenting similar values to Tibetan lambs but significant differences from those of plain Hu lambs. The hypoxia exposure of parents could account for the improved oxygen regulatory ability in their offspring under hypoxia stress.

### Expression of genes associated with high-altitude adaptation and diseases in human

We first examined the similarity of global expression patterns between sheep and human. We retrieved publicly available RNA-Seq data from the human GTEx consortium (v8) and conducted comparative analysis using 17,279 one-to-one orthologous genes in 14 common tissues (i.e., hypothalamus, pituitary, cerebellum, ileum, colon, leukocyte, spleen, heart, muscle, artery, adipose, lung, liver and kidney). The *t-*SNE-based expression clustering among samples clearly recapitulated tissues rather than species (Fig. 7a). Similar results were observed in the hierarchical clustering of tissues based on median gene expression (Fig. 7b).

**Fig. 7.**
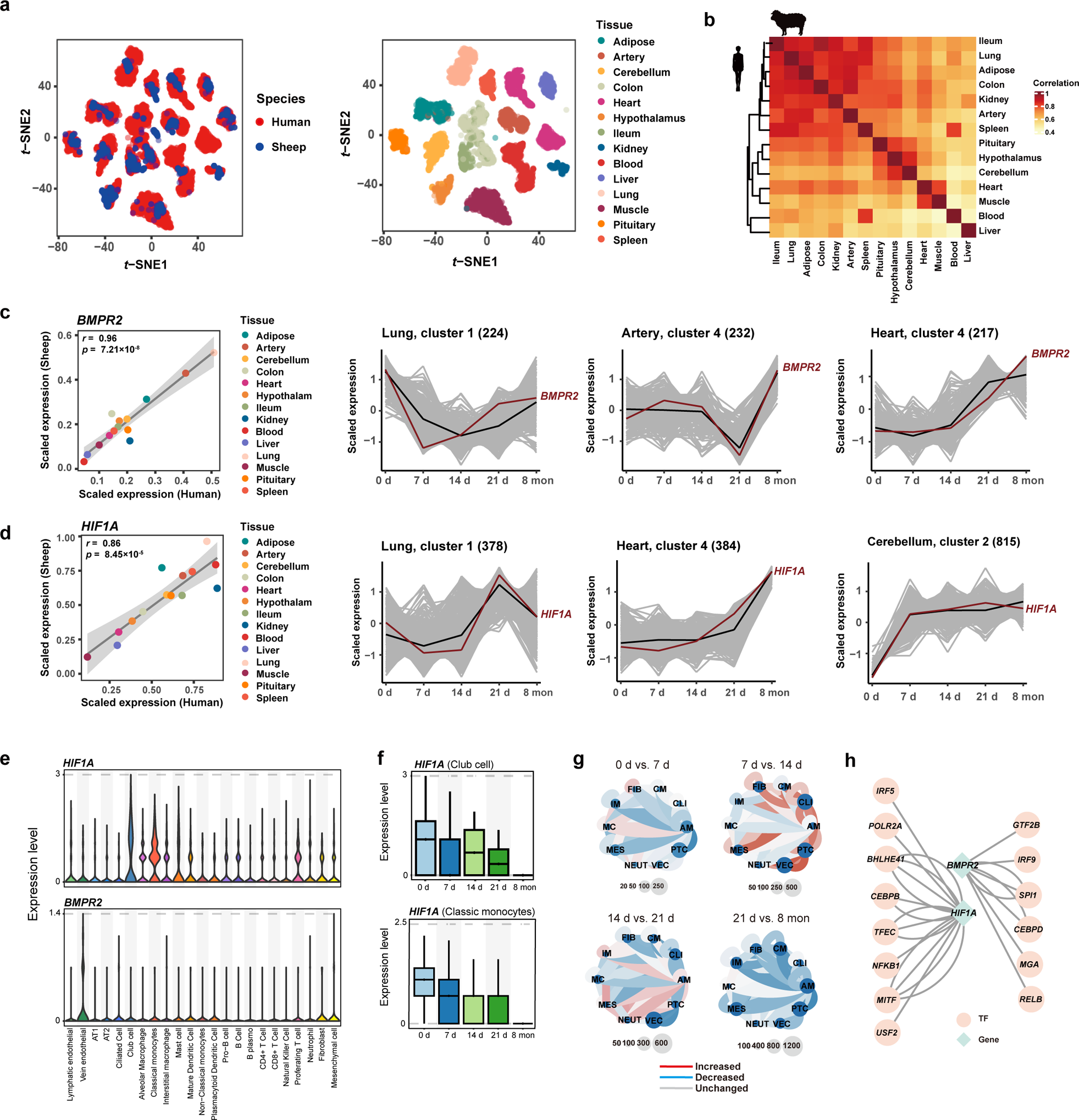
Time-series transcriptome of genes implicated in adaptation and disease in human. a, Conservation of transcripts of 14 common tissues in human and sheep. *t*-SNE clustering of samples in our study (*n* = 1,277) and the human GTEx v8 consortium (*n* = 6,792) based on batch-corrected expression. Species (left) and tissue types (right) are distinguished by colour. b, Hierarchical clustering of common tissues in human and sheep based on Pearson’s correlation of the median TPM value. c-d, Gene examples for human pulmonary hypertension (i.e., *BMPR2*) and high-altitude adaptation (i.e., *HIF1A*). c, Pearson’s correlation between human and sheep based on the median value of *BMPR2* (left). Expression patterns of *BMPR2* in crucial tissues over time (right). d, Similar to c, but for the *HIF1A* gene. e, The expression of *BMPR2* and *HIF1A* across cell types in lung. f, The expression of *BMPR2* in club cells (top) and classic monocytes (bottom) over time in lung. g, Cell-cell communication results for differentially expressed cell types from *BMPR2* and *HIF1A* in adjacent time point comparisons. h, Transcription factors (TFs) regulating *HIF1A* and *BMPR2*.

We collected candidate genes associated with high-altitude adaptation (e.g., Tibetan, Andean and Ethiopians) (Supplementary Table 28) and mountain sickness (e.g., pulmonary hypertension, polycythemia and heart failure) in human (Supplementary Table 29). We calculated the expression correlations of the adaptive and disease-related genes between sheep and human and found that most adaptive genes (97.82%, 1,160 out of 1,207 genes) and disease-related genes (97.43%, 580 out of 613 genes) were significantly correlated (Pearson’s correlation, *P* < 0.05) (Supplementary Table 30). Therefore, we used the sheep time-series transcriptomic data to investigate the expression changes in these genes with time across tissues (Supplementary Figs. 14 and 15). For example, *BMPR2* is a well-known gene associated with pulmonary hypertension^55,56^, and its expression levels were highly correlated (Pearson’s *r* = 0.96, *P* = 7.21×10^−8^) between human and sheep across tissues (Fig. 7c). The expression changes in *BMPR2* in critical tissues such as lung, artery and heart showed distinct patterns over time. Specifically, the expression level changed greatly at 7 d in lung and at 21 d in artery and gradually increased with time in heart (Fig. 7c). Additionally, the expression levels of the high-altitude adaptive gene *HIF1A* displayed a high correlation (Pearson’s *r* = 0.86, *P* = 8.45×10^−5^) between human and sheep (Fig. 7d). The expression of this gene fluctuated with time in lung but gradually increased with time in heart and cerebellum (Fig. 7d). We noted that genes responsible for similar traits or diseases showed similar time-series expression patterns in relevant tissues (Supplementary Table 31). For instance, *APOLD1* and *KCNA5*, which are associated with pulmonary hypertension, showed similar expression patterns to *BMPR2* in lung, artery and heart (Supplementary Table 31). The adaptive genes *EPAS1* and *VEGFA* exhibited similar expression patterns to *HIF1A* in cerebellum and kidney (Extended Data Fig. 15 and Supplementary Table 31).

Furthermore, we used lung scRNA-Seq data to dissect the particular cell types involved in high-altitude adaptation and diseases (Supplementary Tables 32 and 33). We examined the expression of *HIF1A* and *BMPR2* across all cell types in lung tissues. We found high expression levels of *HIF1A* in classical monocytes (CMs) and club cells (CLU) and *BMPR2* in vein endothelial cells (VECs) (Fig. 7e). Additionally, we found that the time-series expression patterns of *HIF1A* and *BMPR2* were similar to those obtained from bulk RNA in lung (Fig. 7f). Cell-cell communication analysis showed that cell communication events continuously increased with time, in which proliferating T cells (PTCs), CMs and CLUs maintained high levels of communication. Moreover, we predicted that core transcription factors (TFs), such as *NFKB1*, *RELB* and *CEBPB*, regulate *HIF1A* and *BMPR2* (Fig. 7h). Notably, *NFKB1*, a transcription regulator of *HIF1A*, can be activated by oxidant-free radicals and ultraviolet irradiation^57^, which is functionally relevant to high-altitude adaptation. The function of the *BMPR2-*related transcription regulator *RELB* is associated with the NF-kappa-B pathway, which is involved in disease-related processes such as inflammation, immunity and tumorigenesis^58^.

## Discussion

Using sheep as a model, we have created the first comprehensive time-series transcriptome atlas of whole-body major tissues for lowland animals transplanted to a high-altitude environment and transcriptomes for their offspring born at high altitudes. Leveraging these critical data, we are able to explore the gene expression patterns of major tissues during the acclimatization process, which offers an exceptional opportunity to dissect the differences in the regulatory mechanisms underlying genetic adaptation and short-term acclimatization to hypoxia as well as the transcriptional changes responsible for the heritance of acclimatization. The sheep time-series transcriptomes also provide valuable resources for depicting the temporal expression of genes associated with human high-altitude adaptation and diseases.

The high-altitude hypoxia adaptation of indigenous highland inhabitants has always been related to the morphological and functional remodelling of the lungs, heart and artery^3,59,60^. However, in the short-term acclimatization of plateau Hu sheep transplanted from lowlands, we found that the cerebellum, kidney, adipose, muscle, colon, ileum, blood and spleen were among the most active tissues in the response to hypoxia stress based on the numbers of DEGs identified between adjacent time points (Fig. 3a) and DCGs across the examined time points (Fig. 3h). This provided clear evidence that different body systems involving distinct tissues and consequently different strategies for oxygen utilization should be required for long-term adaptation and short-term acclimation to hypoxia, respectively. For instance, the nervous (cerebellum), metabolic (kidney, adipose, muscle), digestive (colon, ileum) and immune (blood, spleen) systems seem to make major contributions to short-term hypoxia acclimatization. These systems and tissues may have to reduce oxygen consumption under hypoxia because they are oxygen-consuming parts of the body^61–63^. In contrast, the respiratory (lung) and cardiovascular (heart and artery) systems are mainly responsible for long-term hypoxia adaptation, and they can improve the exchange and transportation of oxygen in response to hypoxia^64,65^. Thus, multiple systems and tissues actively respond to short-term hypoxia acclimatization because short exposure to hypoxia can stimulate the stress response of the whole body, leading to dramatic physical adjustments^30,66^. In this context, we speculate that plateau Hu sheep may cope with a hypoxic environment through the coregulation of central coordination, stress hormones, energy metabolism, intestinal homeostasis and the immune response based on the GO terms of DEGs in relevant tissues (i.e., cerebellum, kidney, adipose, muscle, colon, ileum, blood and spleen) between adjacent times (Supplementary Table 14). The importance of regulating energy metabolism for short-term hypoxia acclimatization is also supported by the functions of tissue-shared DEGs between adjacent times and the functions of DCGs in gene cluster 1 of cerebellum, which were mostly enriched in GO categories of energy metabolism (Fig. 3i). By integrating the expression profiles and bioindicators of blood, we identified 10 gene modules (e.g., M3) that were significantly correlated with informative bioindicators of the hypoxia response, such as erythropoietin (EPO), nitric oxide synthetase (NOS), glutathione peroxidase (GSH-Px) and alkaline phosphatase (ALP) (Fig. 2e). The functions of these bioindicators suggested that the transcriptional regulation of oxygen transportation (EPO, NOS)^67,68^, antioxidation (GSH-Px)^69^ and energy metabolism (ALP)^70^ through the blood and circulatory systems may also contribute to short-term hypoxia acclimatization. Notably, the level of SpO_2,_ one of the most important indicators of blood oxygen content, recovered with acclimatization across time points (Fig. 2a), indicating the successful gradual acclimatization of plateau Hu sheep through the collaborative regulation of the aforementioned different systems and tissues.

Based on the whole-genome selection test between highland and lowland sheep together with differential expression analysis across tissues, we identified 52 *F*_ST_ genes that were differentially expressed between Tibetan sheep and plain Hu sheep, and 179 *F*_ST_ genes that showed dynamic changes in expression during the acclimatization of transplanted Hu sheep in at least five tissues (i.e., multi-tissue *F*_ST_ gene) (Fig. 4a-c). These two panels of multi-tissue *F*_ST_ genes appeared to be putatively selected in the genome and effectively expressed in multiple tissues, providing new clues for delineating the genetic basis of long-term adaptation and short-term acclimation to high-altitude hypoxia at the multi-tissue level. Notably, we found 13 genes (e.g., *APOLD1*, *NDUFB9*, *ERBB4*, *NFKBIZ*, *NR4A3*, *RPS8*, *CIAO2A*, *AHCYL2*, *ESRRG*, *KIAA0930*, *RASGEF1B*, *MRPS25* and *TNFRSF21*) that were shared between the two panels (Fig. 4c), which could represent reliable candidates for both hypoxia adaptation and acclimatization at the genomic and transcriptional levels. For example, *APOLD1* and *NR4A3* showed significant expression changes in many tissues (Fig. 4a, b), and they were reported to be functionally associated with cardiopulmonary, metabolic and neurological phenotypes^71–74^ (Supplementary Table 21). This implied roles of these genes in hypoxia adaptation (cardiopulmonary changes) and acclimation (metabolic and neurological adjustments). Additionally, we explored the roles of CREs in gene expression by integrating differential expression and differential chromatin accessibility analysis. Among the common genes showing both up- or downregulated expression and changes in chromatin accessibility, most genes in plateau Hu sheep and Tibetan sheep were down-regulated compared with those in plain Hu sheep across tissues (Fig. 5f and Supplementary Table 25). The results were consistent with previous findings showing that negative regulatory feedback loops play an important role under hypoxic stress^75,76^. Nevertheless, compared with Tibetan sheep, more common genes in plateau Hu sheep were up-regulated across tissues (Supplementary Table 25), revealing a stronger response to hypoxic conditions for short-term acclimation. In summary, the identified multi-tissue *F*_ST_ genes and common regulated genes can serve as a valuable reference for investigating expression and regulation in different tissues and screening causal variants for hypoxia-related traits.

Interestingly, we found that plateau Hu lambs exhibited SpO_2_ values and gene expression patterns similar to those of Tibetan lambs (Fig. 6a, e), implying that plateau Hu lambs had already acclimatized to high-altitude hypoxia at birth. Notably, the DEGs detected in the comparisons of plateau Hu lambs vs. plain Hu lambs and Tibetan lambs vs. plain Hu lambs were enriched in some common GO terms (e.g., extracellular matrix organization in kidney, localization within membrane in cerebrum and cellular response to angiotensin in artery) that could be directly activated by hypoxia^51^ (Fig. 6c and Supplementary Table 27). The observation suggested similar genetic regulation of relevant tissues (e.g., cerebrum, kidney and artery) in response to hypoxia in plateau Hu lambs and Tibetan lambs. The enriched functions of DEGs also showed GO categories specific to plateau Hu lambs in response to hypoxia in the lungs and artery (Fig. 6d and Supplementary Table 27). This may indicate that plateau Hu lambs descended from short-term hypoxia-acclimatized parents should require more transcriptional regulation and physical adjustments to cope with hypoxia than indigenous Tibetan lambs. Compared with other lowland animals transplanted to high altitudes (e.g., cattle^77^), sheep showed much less neonatal mortality based on both the observations made in our transplantation experiment and previous reports^78^. Since hypoxia may increase stillbirth and infant mortality^79,80^, the DEGs identified in plateau Hu lambs may hold the potential to dissect the genetic basis of low mortality of offspring under hypoxia and thus contribute to the improvement of pregnancy outcomes in human^18^ and other animals.

Our comprehensive transcriptome data for major tissues in the sheep model showed high correlations with human GTEx data according to tissue type, based on the expression clustering of 17,279 one-to-one orthologous genes in 14 common tissues (Fig. 7a, b). This demonstrated that our sheep expression atlas could be used to improve the interpretation of the genetic mechanisms underlying hypoxia-related adaptation and diseases in human. At the gene level, the expression levels of two representative human genes (i.e., *BMPR2*, associated with pulmonary hypertension^55,56^, and *HIF1A*, associated with high-altitude hypoxia adaptation^4,81^) exhibited significantly high correlations in relevant tissues between human and sheep (Fig. 7c, d and Supplementary Table 30), further demonstrating the rationality of employing sheep expression profiles for illustrating the expression patterns of human adaptation and disease genes. Importantly, the time-series characteristics of sheep transcriptomes may be valuable for providing missing information about temporal expression changes in the aforementioned human genes (Fig. 7c, d). Additionally, we created time-series scRNA data for lung tissues, which enabled us to further determine the cell types and cellular mechanisms underlying the expression patterns of human genes (Fig. 7e-h). We expect that our time-series sheep transcriptomes reflect analogous dynamic expression changes in investigated human genes at the tissue and cell levels (e.g., lung).

In conclusion, we generated time-series transcriptome resources of major tissues in a sheep model for high-altitude- or hypoxia-related studies. We identified a credible set of active tissues and crucial genes for the short-term hypoxia acclimatization of sheep and for the inheritance of acclimatization by their offspring. These tissues and genes likely function in multiple body systems and may work together to shape adaptive or maladaptive traits in response to hypoxia. We further utilized sheep time-series transcriptomes to mirror the dynamic expression changes in high-altitude adaptation and disease genes of human, which will probably provide novel insights into the molecular mechanisms underlying human adaptation and diseases. Our study demonstrates how multi-tissue expression profiles across time can be used to inform multiple aspects of short-term acclimatization and disease interpretation.

## Methods

### Experimental design

Individuals of Hu sheep and Tibetan sheep, two representative Chinese native breeds that originally inhabited plains regions (Zhejiang Province, China) and the high-altitude plateau regions [the Qinghai-Tibet Plateau (QTP), China], respectively, were included in the experiment. Overall, adult ewes (∼ 1.5 years) and lambs (∼ 2 months) of the two breeds with good health and body condition were raised under three different scenarios: native on a plain (scenario 1), transplanted from a plain to plateau (scenario 2), and native on a plateau (scenario 3) (Fig. 1a). In scenario 1, 10 adult ewes (∼ 1.5 years) and 6 lambs (∼ 2 months) of Hu sheep were housed at ∼ 350 m.a.s.l. on the Wanghu Livestock Farm in Neijiang City, Sichuan Province, China. In scenario 2, 40 adult ewes and 3 rams of Hu sheep born and raised in the livestock farm under scenario 1 were transplanted to the Tibetan Sheep Breeding Farm of Sichuan Province (Aba Tibetan and Qiang Autonomous Prefecture, Sichuan Province, China) and produced 10 lambs after approximately 8 months. In scenario 3, 10 adult ewes and 6 lambs of Tibetan sheep were housed at ∼ 3,500 m.a.s.l. on the sheep farm under scenario 2. Tibetan sheep have inhabited the QTP for approximately 4,000 years^25,82^. The ewes and lambs were housed and fed similar hay and silage corn, as previously described. and had *ad libitum* access to water and mineral salt.

No statistical methods were used to predetermine the sample size. The experiments were not randomized, and the investigators were not blinded to allocation during experiments and outcome assessment.

### Collection of blood biochemical data and animal tissues

#### Biochemical data and tissue sources

Phenotypic data of 20 blood parameters (i.e., blood oxygen saturation, glucose, triglycerides, alanine transaminase, aspartate aminotransferase, total bilirubin, alkaline phosphatase, lactate dehydrogenase, uric acid, blood urea nitrogen, creatinine, cardiac enzymes, calcium, superoxide dismutase, glutathione peroxidase, catalase, nitric oxide, malondialdehyde, erythropoietin and nitric oxide synthase) and samples from 19 whole-body tissues [i.e., heart, liver, spleen, lung, kidney, rumen, abomasum, duodenum, ileum, jejunum, colon, cerebrum, cerebellum, hypothalamus, pituitary, artery, muscle, adipose and blood (leukocyte)] were collected from all the animals in the above three scenarios. In scenario 2, data and tissues of the 40 ewes of Hu sheep were collected at four sequential time points [7 days, 14 days, 21 days and ∼ 8 months (245 days)] after their transplantation to the QTP, with 10 ewes sampled at each time point. In particular, 10 tissues of the 6 lambs of Hu sheep born on the QTP were collected at an age of ∼2 months, approximately 8 months after transportation (Fig. 1a). In scenarios 1 and 3, the same phenotypic data and tissues of 10 ewes and 6 lambs of Hu sheep (scenario 1) and 10 ewes and 6 lambs of Tibetan sheep (scenario 3) were collected (Supplementary Table 6). Blood oxygen saturation (SpO_2_) values and 19 additional blood biochemical indicators were examined in the animals (Supplementary Table 7). Arterial SpO_2_ was measured using the Tough/Ear Blood Oxygen Metre Veterinary SpO_2_ PR Monitor (RocSea, Jingzhou, China) when the animals were stationary. Blood samples were collected with a 5 mL vacuum tube and were centrifuged immediately to isolate plasma. The isolated plasma samples were then stored in a −80 °C freezer and used to measure the 19 blood biochemical indicators with commercial assay kits (Jincheng Bioengineering Inc., Nanjing, China).

#### RNA-Seq and ATAC-Seq samples

Animals were slaughtered by carotid artery exsanguination. Following sacrifice, tissues were isolated and placed on an ice board for dissection. Each tissue was cut into 5-10 pieces of approximately 50-200 mg each. Samples were then transferred into 2 mL cryogenic vials (Corning, NY, USA, Cat. No. 430917), snap frozen in liquid nitrogen, and stored until RNA extraction for RNA-Seq. In total, 1,277 samples from 19 various tissues of 78 sheep were collected for bulk RNA-Seq (Fig. 1b). Additionally, 66 samples from eight tissues (i.e., hypothalamus, rumen, heart, lung, liver, duodenum, spleen and adipose) of 12 sheep were collected for ATAC-Seq, including 18 samples of three tissues (lung, heart and hypothalamus) from six lambs (two lambs of Hu sheep in the plain, two lambs of Hu sheep in the QTP, and two lambs of Tibetan sheep in the QTP) and 48 samples of eight tissues from six ewes (two ewes of Hu sheep in the plain, two ewes of Hu sheep in the QTP, and two ewes of Tibetan sheep in the QTP) (Fig. 1b).

#### ScRNA-Seq samples

Six samples (i.e., plain Hu sheep in scenario 1, plateau Hu sheep at four acclimatization time points in scenario 2 and Tibetan sheep in scenario 3) from different parts of the lung (left lung and right lung) were harvested and then cleaned with PBS for scRNA-Seq. For each sample, sliced tissues were stored in tissue storage solution (Miltenyi Biotec, Bergisch Gladbach, Germany, Cat. No. 130-100-008) at 4 °C for single-cell suspension preparation and library construction.

### RNA extraction, library preparation and sequencing

Total RNA was extracted from flash-frozen tissues with RNA TRIzol (Invitrogen, Carlsbad, CA, USA) according to the manufacturer’s protocol. After purification, RNA quality was checked using agarose gel electrophoresis and a NanoPhotometer® spectrophotometer (IMPLEN, CA, USA). RNA integrity (RIN) was examined on an Agilent 2100 Bioanalyzer (Agilent Technologies, Waldbronn, Germany) with a cut-off of an RIN < 7.00, and the RNA concentration was measured with the Agilent 2100 RNA 6000 Nano Kit (Agilent Technologies, Waldbronn, Germany). First-strand cDNA was generated using the FastKing One-Step RTLPCR Kit (TIANGEN Biotech, Beijing, China), and cDNA libraries were constructed by the Illumina TruSeq RNA Library Prep Kit v2 (Illumina, CA, USA). RNA-Seq was implemented on the Illumina HiSeq 2500 (Illumina, CA, USA) at Novegene Co., Ltd. (TianJin, China), generating 150 bp paired-end reads.

### ATAC library construction and sequencing

Library preparation for ATAC-Seq followed a modified OmniATAC protocol^83^ in cryopreserved nuclei. Specifically, weighed frozen tissues (∼ 20 mg) were first lysed in cold homogenization buffer (10 mM Tris-HCl, pH 7.4, 10 mM NaCl, 3 mM MgCl_2_, 0.1% Igepal). Nuclei were then resuspended and collected from the interface after iodixanol-based density gradient centrifugation. Thereafter, nucleus tagmentation was performed in Tn5 transposase reaction mix (Illumina Tagment DNA Enzyme and Buffer kits) under incubation at 37 °C for 30 min in a thermomixer with shaking at 1,000 rpm, and two equimolar adapters were added. Immediately following the transposition reactions, DNA was purified with the Qiagen MinElute PCR Purification Kit (Qiagen, Netherlands, Cat. No. 28004) and eluted in EB buffer. To amplify the library, PCR was then performed in a mix of 10 µM Nextera i7 and i5 primers and NEBNext Q5 High-Fidelity PCR Master Mix (New England Biolabs, MA, USA) according to the following protocol: 72 °C for 5 min, 98 °C for 30 sec, and 11 cycles of 98 °C for 10 sec, 63 °C for 30 sec and 72 °C for 1 min. PCR products were purified with the Qiagen MinElute PCR Purification Kit and AMPure XP beads (Beckman Coulter, Cat. No. A63880) and resuspended in ultrapure nuclease-free distilled water. Library quality was assessed with a Qubit 2.0 system (Life Technologies, MA, USA), and fragment size was examined using an Agilent 2100 Bioanalyzer. The libraries were sequenced on an Illumina NovaSeq 6000 system with a 150 bp paired-end sequencing method.

### scRNA-Seq library construction and sequencing

scRNA-Seq libraries of lung tissues were constructed following previous protocols with minor modifications ^84–86^. For the lung tissues, gentle and rapid generation of single-cell suspensions was achieved by using a modified version of the procedure of a mouse Lung Dissociation Kit (Miltenyi Biotec, Bergisch Gladbach, Germany; Cat. No. 130-095-927). In summary, we dissected sheep lung tissue into single lobes and rinsed the lobes in petri dishes containing PBS (pH = 7.2) to remove residual vessels, blood clots and mucin. Clean lobes were subjected to shacking digestion at 37 °C for 25-30 min with enzyme mix, which consisted of 2.4 mL of 1× Buffer S, 100 µL of Enzyme D, and 15 µL of Enzyme A. The digestion solution was briefly centrifuged at 600 × g for 2 min at 4 °C, and the precipitated pellet was resuspended in 2.5 mL 1× Buffer and filtered with a 70 µM MACS SmartStrainer (Miltenyi Biotec, Bergisch Gladbach, Germany; Cat. No. 130-098-462). Then, the obtained cell suspension was centrifuged at 300 × g for 10 min at 4 °C. After removing the supernatant, the cell pellet was resuspended in an appropriate buffer to the required volume for scRNA-Seq.

Qualified single-cell suspensions containing at least 8,000 cells were loaded onto a chromium single-cell controller (10× Genomics), and single-cell gel beads were generated in the emulsion according to the manufacturer’s protocol. Then, scRNA-Seq libraries were constructed using Single Cell 3’ Library and Gel Bead Kit v3.1 (8,000 initial cell capture number) and were subsequently sequenced using a NovaSeq 6000 sequencer (Illumina).

### Whole-genome sequence (WGS) data

#### Data collection

Whole-genome sequences of 37 sheep (average depth = ∼ 17×) were retrieved from five previous studies^25,26,87–89^, including 17 sheep of 2 populations from the high-altitude region (∼ 3,700 m.a.s.l.), and 20 sheep of 5 populations from the low-altitude region (∼ 17 m.a.s.l.). Detailed information on the populations, including the names, sampling locations and number of samples, is summarized in Supplementary Table 8.

#### Variant calling

SNP calling followed previous protocols^26^. First, we filtered low-quality bases and artefact sequences using Trimmomatic (v.0.36)^90^ and aligned the high-quality paired-end reads (150 bp or 100 bp) to the sheep reference genome *Oar_rambouillet_v1.0*. (GCA_002742125.1) using BWA (v0.7.8)^91^ with the default parameters. Next, we removed duplicates in the BAM files using the *MarkDuplicates* module in GATK (v4.1.2.0)^92^. SNPs were then detected using the GATK *HaplotypeCaller* module with the GATK best-practice recommendations. Thereafter, we merged the GVCFs files called individually by the *CombineGVCFs* module and called SNPs with the *GenotypeGVCFs* module. Finally, we selected the raw SNPs using the *SelectVariants* module and filtered them using *VariantFiltering* of GATK with the parameters “QUAL < 30.0 || QD < 2.0 || MQ < 40.0 || FS > 60.0 || SOR > 3.0 || MQRankSum < −12.5 || ReadPosRankSum < −20.0”.

#### SNP quality control

SNP quality control was conducted with the following criteria using VCFtools (v0.1.17)^93^: 1) call rate > 90% and 2) minor allele frequency (MAF) > 0.05. Any SNPs that failed to meet any of the above criteria were filtered, and we obtained 34,355,535 SNPs for downstream analysis.

### Hi-C data preprocessing

#### Quality control and data preprocessing

Hi-C data from the blood of sheep were retrieved from the NCBI Sequence Read Archive (SRA) under accession number SRR19426890. We first trimmed adapter sequences and low-quality reads with Trimmomatic (v.0.36)^90^ and obtained ∼ 780 million clean reads. Hic-Pro (v2.9.0)^94^ was then used to process the Hi-C data from raw sequencing data via a pipeline including alignment, matrix construction, matrix balancing, and iterative correction and eigenvector decomposition (ICE) normalization with the default parameters.

#### Detection of TADs

To explore the ATAC-Seq peak-to-gene linkage, we identified topologically associating domains (TADs) as follows. We first implemented the conversion of Hi-C matrices to Cooler format via HiCPeaks (v0.3.2)^95^. To detect the TADs, we then calculated the directionality index (DI) with a resolution of a Hi-C matrix at 40 kb using the *hitad* function from TADLib (v0.4.2)^96^. In total, we obtained 3,032 TADs for subsequent analysis.

### RNA-Seq data preprocessing

Raw RNA-Seq reads with low base quality scores (quality scores ≤ 20) were first trimmed, and then adapter contamination was further removed using fastp (v0.20.1)^97^. The high-quality clean reads were next mapped to the sheep reference genome *Oar_rambouillet_v1.0* (GCA_002742125.1) by the program STAR (v2.7.9a)^98^ with the settings “-quantMode GeneCounts, -outFilterMismatchNmax 3, -outFilterMultimapNmax 10”. Properly paired and uniquely mapped reads were extracted by using SAMtools (v1.11)^99^ with the command “view -f 2”. Gene counts were generated by the featureCounts program from the Subread package suite (v2.0.3)^100^. We also normalized the raw counts of genes using the transcripts per million (TPM) method with an in-house script.

### ATAC-Seq data preprocessing

#### Mapping and peak calling

Raw ATAC-Seq reads were trimmed for Nextera adaptors by using Fastp (v0.20.0)^97^ with the options “-q 15 -l 18”, and the clean reads were aligned to the sheep reference genome *Oar_rambouillet_v1.0* using BWA (v0.7.17)^91^ with the default parameters. PCR duplicates were removed using Picard (v1.27)^92^, and uniquely mapped high-quality reads were collected using SAMtools (v1.11)^99^ with the options “view -f 2 -q 30”. The reads mapped to the mitochondrial genome were also discarded, and the final BAM files were kept for subsequent analysis.

ATAC-Seq peaks were called by Genrich (v0.6.1)^101^ with the options “-j -p 0.01 -b”. Peak calling was first implemented in each library and then for each tissue based on concatenating all the replicates by using BEDTools (v2.30.0)^102^ with the *bedtools merge* function. The *P* value was calculated for each peak assuming a null model with a log-normal distribution and corrected based on the BenjaminiLHochberg model.

#### Quality control and data reproducibility

The following parameters were measured as recommended by the ENCODE project (https://www.encodeproject.org/atac-seq/#standards) for the validation of ATAC-Seq libraries. The fraction of reads in peaks (FRiP) scores, nonredundant fraction (NRF) and other quality metrics (e.g., PCR bottlenecking coefficients, PBCs) for each sample were calculated with either SAMtools (v1.11)^99^ or in-house scripts. Details of the quality control scores are included in Supplementary Table 4. Global correlations between samples were calculated with the R package DiffBind (v3.2.7)^103^.

#### Consensus ATAC-Seq peaks

To generate consensus peaks, peaks for individual samples were merged with the “*multiBigwigSummary BED-file*” function in deepTools (v3.5.0)^104^. BAM files were converted into a normalized coverage track of bigWig format using the *bamCoverage* command in deepTools with the options “--binSize 10 --normailzeUsingRPKM -centerReads”, and ATAC peaks were then visualized with Integrative Genomics Viewer (IGV) software (v2.9.4)^105^. Raw read counts in these peaks were determined by “*multiBamSummary BED-file*” and were normalized for reads per kilobase per million (RPKM) reads with the function “*normalize.quantiles*” in the R package preprocessCore (v1.40.0)^106^. Additionally, *t*-SNE clustering of the ATAC-Seq profile was performed as described for the *t*-SNE analysis of the RNA-Seq data above.

### Whole-genome selective sweep tests

To identify potential selective signatures associated with hypoxia adaptation between populations in high- and low-altitude regions, we selected domestic sheep from plateau (*n* = 17) and plain (*n* = 20) areas (Supplementary Table 8). We calculated genome-wide pairwise *F*_ST_ values^107^ between the high- and low-altitude populations with a sliding window approach (10 kb sliding windows with 10 kb steps) using the Python script *popgenWindows.py* (https://github.com/simonhmartin/genomics_general). Regions with the top 5% of the average *F*_ST_ distribution were defined as selective signatures and then annotated to corresponding gained genes.

### Tissue specificity of gene expression and chromatin accessibility

First, we clustered all 1,277 samples based on TPM and the *t*-distributed stochastic neighbour embedding (*t*-SNE) method as implemented in the R package Rstne (v0.16)^108^. To examine tissue similarity, median gene expression for each tissue was calculated and then used to cluster tissues based on the Euclidean distance with the corresponding function in the R package ComplexHeatmap (v2.8.0)^109^.

Then, the 19 whole-body tissues were classified into 10 different broad categories (i.e., muscle, immune, intestine, rumen, brain, artery, adipose, liver, lung and kidney) (Supplementary Table 1). To characterize the genes showing tissue-specific expression, we compared gene expression in the samples of a target tissue with that in tissues from the other categories^110^ using the R package limma (v3.48.3)^111^. Known covariates (e.g., age and time point) were calibrated via the *combat* function of the R package sva (v.3.40)^112^. We ranked *t-*statistics computed by limma and considered the top 5% of genes with an absolute value of log_2_-transformed fold-change (|log_2_(FC)|) > 1 and a false discovery rate (FDR) < 0.01 as tissue-specific genes (TSGs).

For the ATAC-Seq data, we used a Shannon entropy-based method^113^ to compute the tissue specificity index with a normalized peak matrix. Specifically, for each peak, we defined its relative accessibility in tissue *i* as 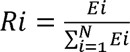, where *E*_i_ is the normalized median reads per kilobase million (RPKM) value for the peak in tissue *i*, 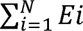 is the sum of normalized median RPKM values in all tissues, and *N* is the total number of tissues. The peak entropy score across tissues can be defined as 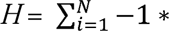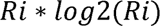, where the value of *H* ranges between 0 and log_2_(*N*). An entropy score close to zero indicates highly tissue-specific accessibility of the peak; conversely, a score close to log_2_(*N*) indicates ubiquitous accessibility of the peak. We selected peaks with an entropy score < 2.5 as the tissue-restricted peaks, while peaks were assigned to particular tissues with maximal RPKM values. Peaks with the top 500 entropy scores were considered conserved peaks.

### Comparative transcriptome analysis

Human RNA-Seq data normalized by the transcripts per million (TPM) method was generated by the human GTEx (Genotype-Tissue Expression) consortium. From the GTEx v8 release (https://gtexportal.org/home/datasets), we obtained a subset of 6,792 RNA-Seq samples from 14 common tissues and 17,279 one-to-one orthologous genes between human and sheep. We then used the function *IntegrateData* with the parameters “anchorset = expression, dims = 1:30” in the R package Seurat (v4.2.0)^114^ to combine the expression values of orthologous genes in human and sheep by removing hidden confounding factors. Thereafter, we performed *t*-SNE clustering of the samples using a two-dimensional projection based on corrected expression values of orthologous genes. We calculated the median values of gene expression in each tissue of sheep and human separately, representing the expression of the particular tissue in each species^115^. Additionally, we performed hierarchical clustering using the R package ComplexHeatmap (v2.8.0)^109^ to explore the relationships of tissues based on the median values.

### Characterization of gene expression changes along a timeline

In each tissue of the Hu sheep transported to high altitude, we identified significantly differentially expressed genes (DEGs) between adjacent time points (0 d vs. 7 d, 7 d vs. 14 d, 14 d vs. 21 d, and 21 d vs. ∼ 8 mon) with thresholds of an FDR < 0.05 and |log_2_(FC)| > 0.5 using the R package DESeq2 (v1.32.0)^116^. Between the adjacent time points, *k*-means clustering was used to characterize the patterns of gene expression changes in different tissues. We first built a log_2_(FC) matrix and determined the *k* value with the *fviz_nbclust* function in the R package factoextra (v1.0.7)^117^. Common genes among different tissues were then clustered into a set of *k* groups based on the Euclidean distance with the *kmeans* function in R.

Additionally, we collected candidate genes related to high-altitude adaptation and mountain sickness in human from previous studies (Supplementary Tables 28 and 29). Furthermore, we characterized the time-series patterns of gene expression changes across tissues using the same methods described above.

### Time series analysis

Furthermore, we performed time series analysis of gene expression to identify dynamically changing genes (DCGs) over time using the R package maSigPro (v1.64.8)^118^. First, we calculated the raw count matrix and retained genes with a sum of all counts ≥ 10 reads in all the samples. We then performed normalization with the *vst* function in the R package DESeq2 (v1.32.0)^116^. We ran maSigPro with a degree = 4 and considered genes with a goodness-of-fit (*R*^2^) ≥ 0.4 as DCGs. We also applied a soft-clustering approach (*c*-means clustering) in the R package mFuzz (v.2.23.0)^119^ to identify the most common profiles for individual tissues. We designated clusters when increasing the number would not add a new cluster but instead split a previous cluster.

### Weighted gene co-expression network analysis (WGCNA)

We employed the R package WGCNA (v1.12.0)^120^ to construct a gene co-expression network for an aggregated expression matrix in blood. Similar to the time-series analysis, we kept genes with a sum of all counts ≥ 10 reads in all the samples and a median absolute deviation (MAD) value > 0.01 (top 75% of MAD) and then performed normalization with the *vst* function in the R package DESeq2 (v1.32.0)^116^, resulting in 13,707 genes for weighted gene co-expression network analysis. Subsequently, we transformed the normalized matrix to a similarity matrix based on the pairwise Pearson’s correlation among the 13,707 genes and then converted the similarity matrix into an adjacency matrix. By using the dynamic hybrid cutting method, we clustered genes with similar expression patterns (*r* > 0.85) into 14 distinct gene modules and used principal component analysis (PCA) to summarize modules of gene expression with the *blockwiseModules* function. We then used module eigengene values of the first principal component to test the correlation between module expression and phenotypic data.

### Genomic annotation of ATAC-Seq peaks

We annotated the ATAC-Seq peaks using the *annotatePeak* function in the R package ChIPseeker (v1.28.3)^121^. Based on the distance from the peak centre to the transcription start sites (TSS), the peaks were assigned to functional genomic regions such as promoter (TSS ± 2 kb), downstream, distal intergenic, intron, exon, 5′ untranslated region (UTR), and 3′ UTR. We then calculated the percentage of peaks located in the upstream and downstream regions from the TSS of the nearest genes and visualized the distribution using the *plotDistToTSS* function in ChIPseeker.

### Differential accessibility analysis

In each tissue, we defined differentially accessible regions (DARs) among groups (e.g., three groups in dataset 1: the ewes of Hu sheep, the Hu sheep transplanted to the QTP after 8 months and Tibetan sheep; three groups in dataset 2: the lambs of Hu sheep, Hu sheep raised on the QTP for ∼ 8 months and Tibetan sheep) using the *dba.count, dba.contrast, dba.analyze* and *dba.report* functions of the R package DiffBind (v3.6.1)^103^. We set the thresholds as follows: *P* value < 0.05 and |log_2_(FC)| > 0.5.

### Motif enrichment analysis

We converted the locations of target peaks from the sheep reference genome *Oar_rambouillet_v1.0* (GCA_002742125.1) to the human reference genome *GRCh38* (GCA_000001405.15) using the program LiftOver (v377) with the default settings. Then, enrichment analysis of known binding motifs in peaks was performed using the “findMotifsGenome.pl” script in the software HOMER (v4.8)^122^. *P* values were calculated with a hypergeometric test, and a *P* value < 0.05 was regarded as the threshold for identifying significant motifs.

### Gene-linked candidate *cis*-regulatory elements within TAD

We utilized the correlation approach to identify putative causal relationships between ATAC-Seq peaks and gene expression in the same samples^122–124^. For each TAD in the sheep genome defined above, we computed the Pearson correlation coefficient (PCC) between the estimates of chromatic accessibility [log_2_(RPKM+1)] and gene expression [log_2_(TPM+1)]. To examine the significance level (*P* value) of these correlations, a null distribution was estimated empirically by calculating the PCC of the TAD-constrained peaks with all the genes on the chromosome. Significant associations between TAD-constrained peaks and the expression of relevant genes were identified according to a *P* value < 0.05 and PCC ≥ 0.25.

### Functional enrichment analysis

Human homologues of the sheep Ensembl genes were fetched using the R package biomaRt (v2.52.0)^125^. Gene Ontology (GO) enrichment analysis was implemented based on the human database org.Hs.eg.db (v3.16)^126^ to identify significant biological functions using the R package ClusterProfiler (v4.0.5)^127^, with the parameters of OrgDB = org.Hs.eg.db, fun = “enrichGO”, ont = “BP”, pvalueCutoff = 0.05, and pAdjustMethod = “BH”.

### Phenome-wide association analysis (PheWAS)

The GWAS ATLAS database (https://atlas.ctglab.nl/) provides abundant resources for associating a given gene or SNP with a wide variety of human phenotypes; this strategy has been widely used and proven to be a useful approach, complementary to regular GWAS^128,129^. To explore whether human orthologues of key candidate genes detected in the context of hypoxia acclimatization in sheep are associated with similar adaptive traits in human, we performed PheWAS analysis for human orthologous genes across 3,302 human phenotypes. We included only human GWASs with a sample size > 10,000 in the analysis and considered genes with an FDR < 0.05 to be significantly associated with the corresponding phenotypic trait.

### Analysis of phenotypic measures

For the SpO_2_ and 19 additional blood biochemical indices, phenotypic measurement data were represented as the mean ± standard deviation (SD). Statistical analysis was implemented in RStudio v4.2.0. First, we assessed the normality of the dataset with the *shapiro.test* function. Then, statistically significant differences among groups of animals at different time points were determined by one-way ANOVA (analysis of variance) with the function *aov* when the datasets conformed to the normal distribution, and Tukey’s honestly significant difference (HSD) test was performed to correct for multiple comparisons using the *TukeyHSD* function. For the datasets not under a normal distribution, the KruskalLWallis test was used to estimate the significant differences among groups with the function *kruskal.test*, while the Wilcoxon rank test was used to determine differences between pairwise groups with the *pairwise.wilcox.test* function. Variations of each phenotypic indicator were considered statistically significant with a BenjaminiLHochberg adjusted *P* value < 0.05.

### Analysis of scRNA-Seq data

After sequencing, the high-quality reads were aligned to the sheep reference genome *ARS-UI_Ramb_v2.0* (GCA_016772045.1). We created the premRNA reference following the protocol of 10× Genomics and earlier studies^130^. To obtain the filtered count matrix for subsequent analysis, we calculated gene counts using Cell Ranger (v7.0) (https://github.com/10XGenomics/cellranger) with the default parameters.

The single-cell expression matrices obtained as described above were processed following the protocols of the R package Seurat (v.4.1.1)^114^. We retained cells with more than 200 genes and a low percentage (< 30%) of UMIs mapped to mitochondrial genes. We also used the R package DoubletFinder (v.2.0.3)^131^ to mitigate potential doublets with the *doubletFinder_v3* function. We applied the canonical correlation analysis (CCA) method for dataset integration to correct batch effects as follows. First, normalization for each sample was implemented with the *SCTransfrom* function. We then determined features and anchors for data integration using the *PrepSCTIntegration* and *FindIntegrationAnchors* functions with the default parameters. Furthermore, we generated an integrated expression matrix for each tissue via the *IntegrateData* function.

We scaled the integrated data and applied the *RunPCA* function in the dimensional reduction. Cells were clustered with the *FindClusters* function and visualized using uniform manifold approximation and projection (UMAP). Gene markers for each cell type or subtype were found using *FindAllMarkers* with the Wilcoxon rank sum test (*P* < 0.05, |log_2_FC| > 0.25), including known canonical marker genes and novel ones (Supplementary Table 33).

We predicted core regulatory transcription factor (TF)-gene pairs from the scRNA-Seq data using the R package GENIE3 (v1.6.0)^132^ with the RcisTarget database v1.4.0 (https://resources.aertslab.org/cistarget/) as a reference. Specifically, we used GENIE3 and the workflow in R package SCENIC (v1.1.2.2)^133^ with the default parameters to infer TF-target gene regulatory networks based on the gene expression matrices of each tissue and the DEGs of each cell type. Then, RcisTarget was used to identify enriched TF-binding motifs and predict candidate target genes (regulons) based on the hg38 RcisTarget database, which contains cross-species genome-wide rankings for each motif. Only the TF target genes with high-confidence annotations and corresponding transcription regulatory networks were visualized with Cytoscape (v3.7.1)^134^.

CellLcell communication was inferred from the scRNA-Seq data using CellPhoneDB software (v2.0)^132^. Only receptors and ligands expressed in at least 10% of cells of a given cell type were retained for further analysis, whereas the interaction was considered nonexistent if either the ligand or the receptor was unqualified. The average expression of each ligandLreceptor pair was compared among different cell types to identify their cell-specific expression. Only the ligandLreceptor pairs with significant means (*P* < 0.05) were used for subsequent inference of cellLcell communications in each tissue across time points. The visualization of the changes in the number of ligand-receptor pairs along the examined timescale for each tissue was implemented using Cytoscape (v3.7.1)^134^.

### Ethical statement

All experimental protocols in this study were reviewed and approved by the Institutional Animal Care and Use Committee of China Agricultural University (CAU20160628-2) and the local animal research ethics committee. Animal care, maintenance, procedures, and experimentation were performed in strict accordance with the guidelines and regulations approved by the Welfare and Ethics Committee of the Chinese Association for Laboratory Animal Sciences.

## Data availability

All raw data analysed in this study are publicly available for download without restriction from the Sequence Read Archive (SRA) database in NCBI under accession number PRJNA1000743 and PRJNA1001016 for RNA-Seq and PRJNA1001505 for ATAC-Seq and scRNA-Seq data.

## Acknowledgements

This study was financially supported by grants from the National Key Research and Development Program of China (Nos. 2022YFE0113300, 2021YFD1300904, 2021YFD1200901 and 2021YFF1000703), the National Natural Science Foundation of China (Nos. 32272845, 32061133010, 31972527 and U21A20246), and the Initiation Fund of Sanya Institute of China Agricultural University (No. SYND-2022-12). We also thank the High-performance Computing Platform of China Agricultural University.

## Author Contributions

M.H.L. conceived and supervised the study. M.H.L. and Z.Y. designed the project proposal. Z.Y. performed major bioinformatic analysis of multi-omics data. F.H.L. conducted whole-genome sequencing data analysis and W.T.W. performed single-cell RNA-Seq data analysis. Z.Y., M.L.Z., Y.J.L., D.X.M., W.T.W., R.M., X.W., M.M.W. and J.H.H. contributed to the sample collection and resource generation. Z.Y., J.Y. and M.H.L. wrote and revised the manuscript. All authors reviewed, edited and approved the final manuscript.

