## Supplementary tables 1-33 for "A time-resolved multi-omics atlas of transcriptional regulation in response to high-altitude hypoxia across whole-body tissues"

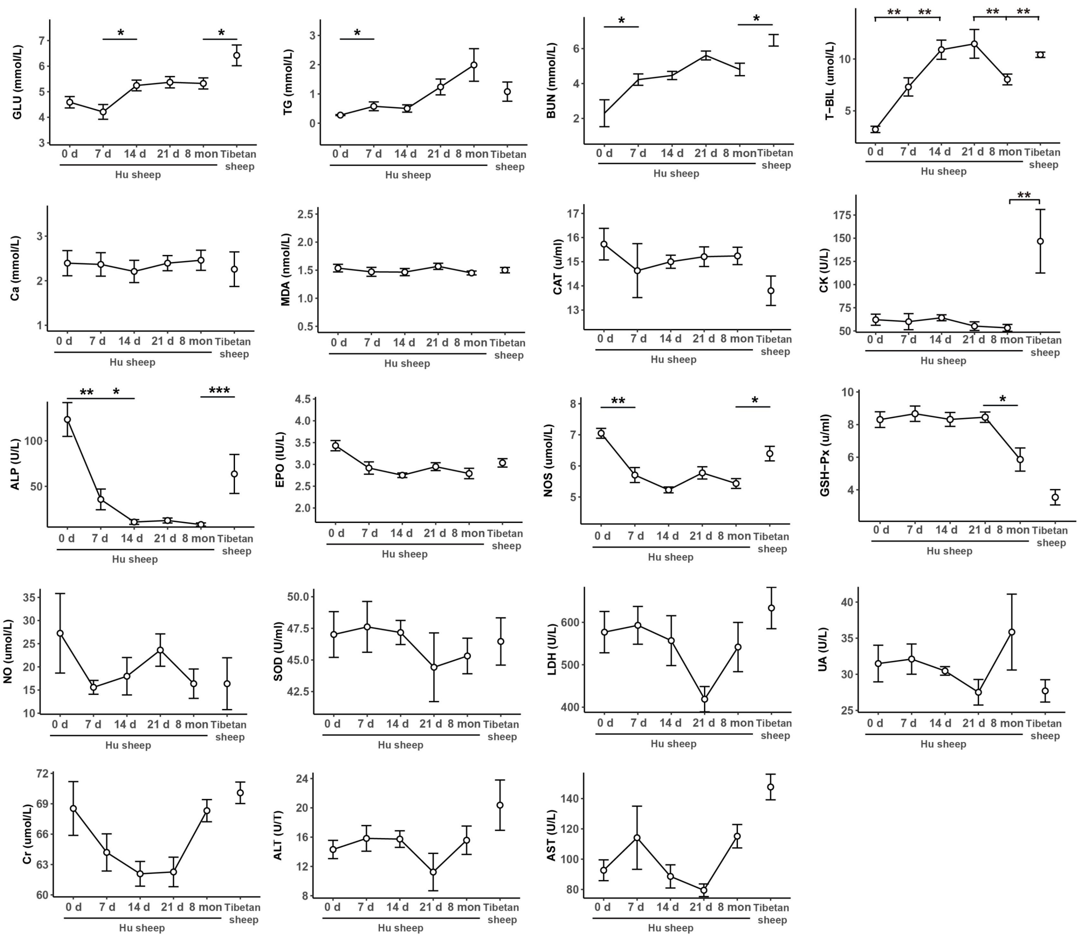


**Extended Data Fig. 1 The changes of phenotypes.** The change in value of bio-indicators with time. *P*-values from Wilcoxon rank sum test, * $P < 0.05$, ** $P < 0.001$, *** $P < 0.0001$.

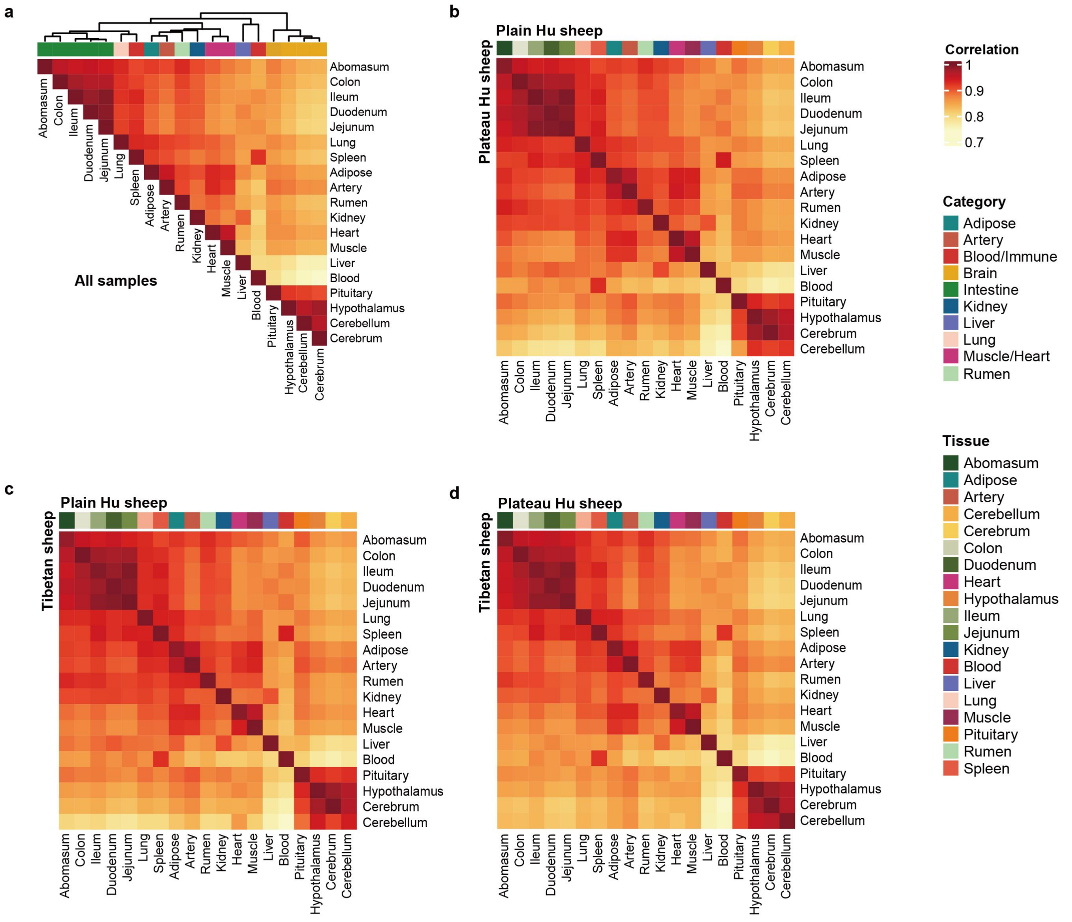


**Extended Data Fig. 2 Hierarchical clustering of RNA-Seq samples.** a, Hierarchical clustering of tissues (*n* = 1,277) based on Pearson's correlation of median value of expression. b-d, Similar with (a), hierarchical clustering of tissues based on Pearson's correlation of median value of expression between plain Hu sheep and plateau Hu sheep (b), between plain Hu sheep and Tibetan sheep (c) and between plateau Hu sheep and Tibetan sheep (d).

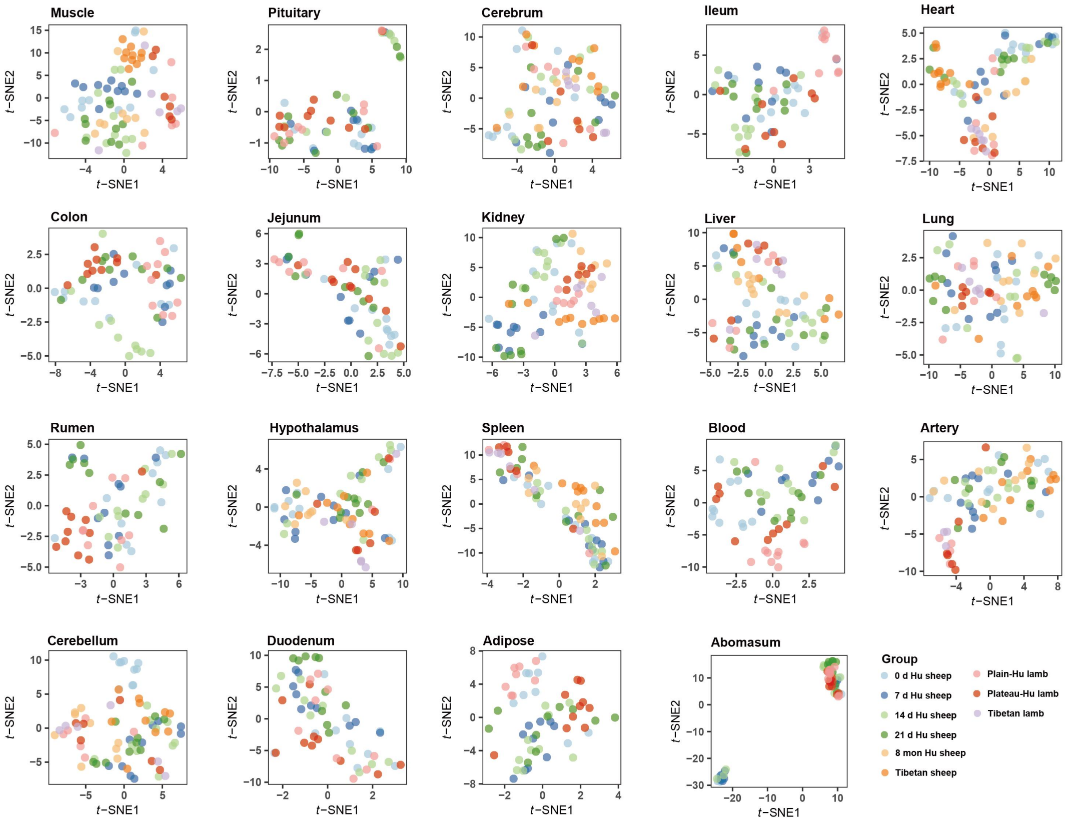


**Extended Data Fig. 3 Sample clustering within each tissue.** *t*-Distributed Stochastic Neighbor Embedding (*t*-SNE) clustering between groups within each tissue.

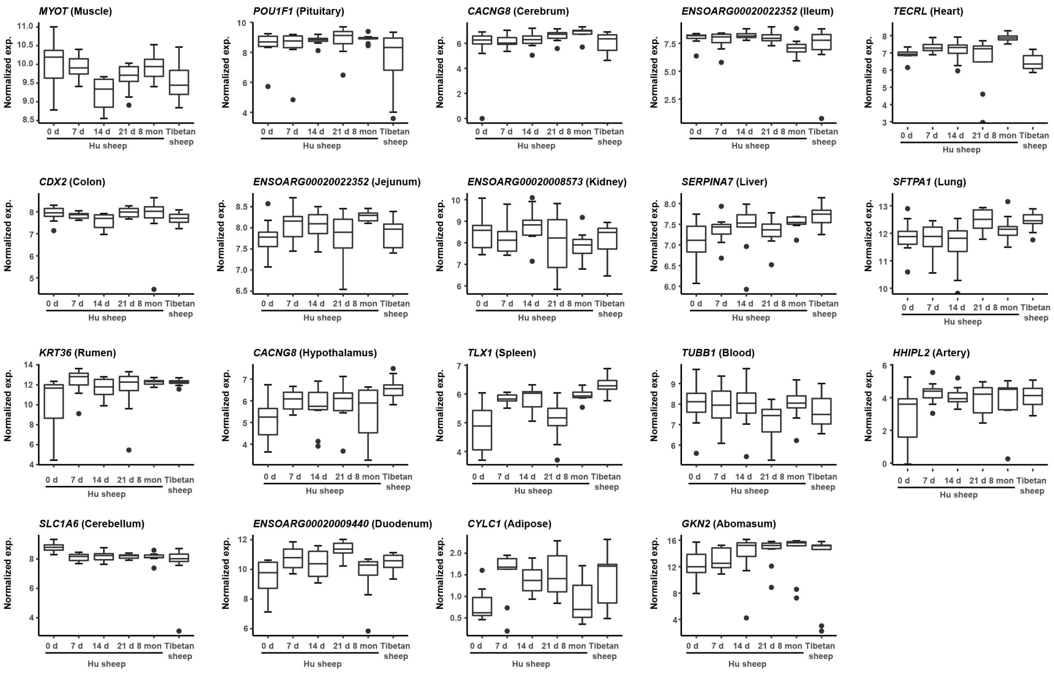


**Extended Data Fig. 4 Example of tissue-specific expressed gene.** The expression level of tissue-specific gene with time in corresponding tissue. Gene with top *t*-statistic is shown for each tissue.

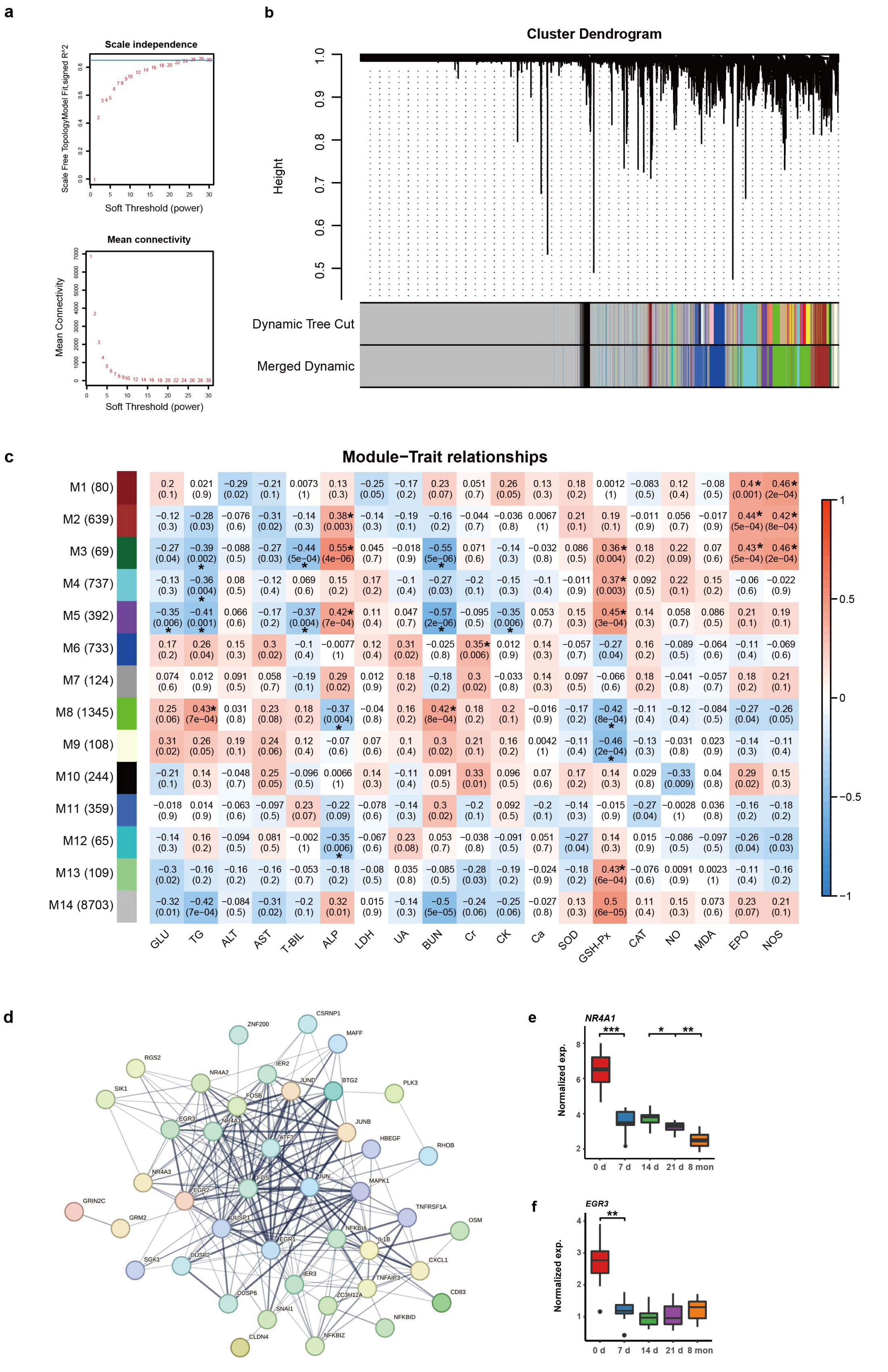


**Extended Data Fig. 5 Weighted correlation network analysis (WGCNA).** a, Identification of optimal β-value. b, The weighted gene co-expression network is constructed in blood. Colors represent gene co-expression modules. c, 14 gene modules (M1-M14) associated with 19 blood bio-indicators (glucose, GLU; triglycerides, TG; alanine transaminase, ALT; aspartate aminotransferase, AST; total bilirubin, T-BIL; alkaline phosphatase, ALP; lactate dehydrogenase, LDH; uric acid, UA; blood urea nitrogen, BUN; creatinine, Cr; cardiac enzymes, CK; calcium, Ca; superoxide dismutase, SOD; glutathione peroxidase, GSH-Px; catalase, CAT; nitric oxide, NO; malondialdehyde, MAD; erythropoietin, EPO and nitric oxide synthase, NOS). The statistical significance of module-trait relationship is corrected for multiple testing using the FDR method, where "*" means FDR < 0.05. The values in the brackets are the numbers of genes in corresponding modules. d, Protein-protein interaction network analysis (STRING database v11) for gene module 3. e-f, Gene examples of gene module 3. The expression changes of *NR4A1* (top) and *EGR3* (bottom) with time. *P* values from Wilcoxon rank sum test, * $P < 0.05$, ** $P < 0.01$, *** $P < 0.001$.

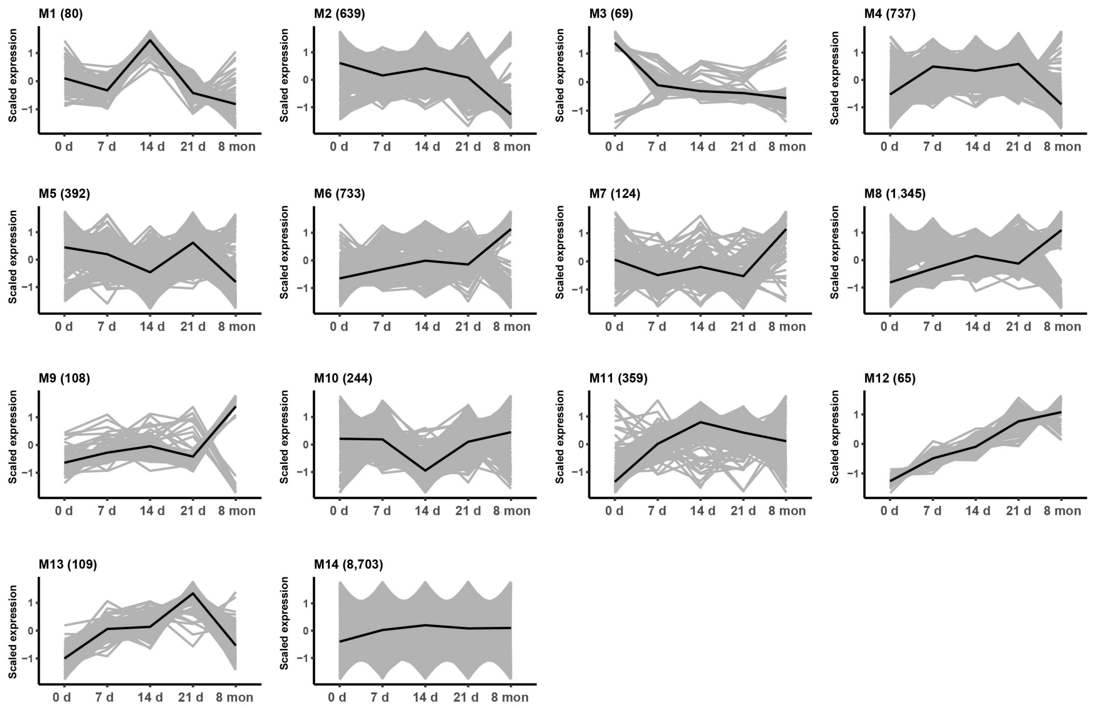


**Extended Data Fig. 6 Time-series expression of 14 gene modules from WGCNA.** The expression changes of 14 gene modules (M1-M14) with time.

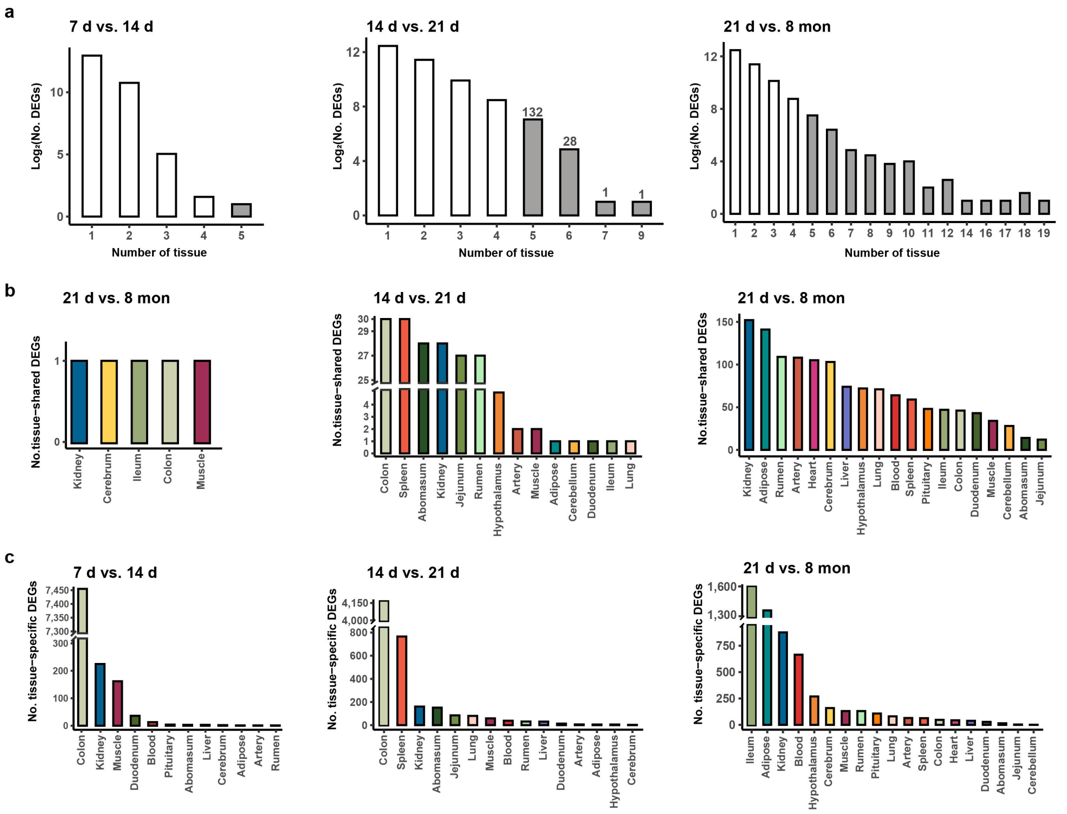

**Extended Data Fig. 7 The distribution of tissue-shared and tissue-specific DEGs.** a, The distribution of DEGs across the number of tissues between different comparisons. b, The number of tissue-shared DEGs across tissues between different comparisons. c, The number of tissue-specific DEGs across tissues between different comparisons.

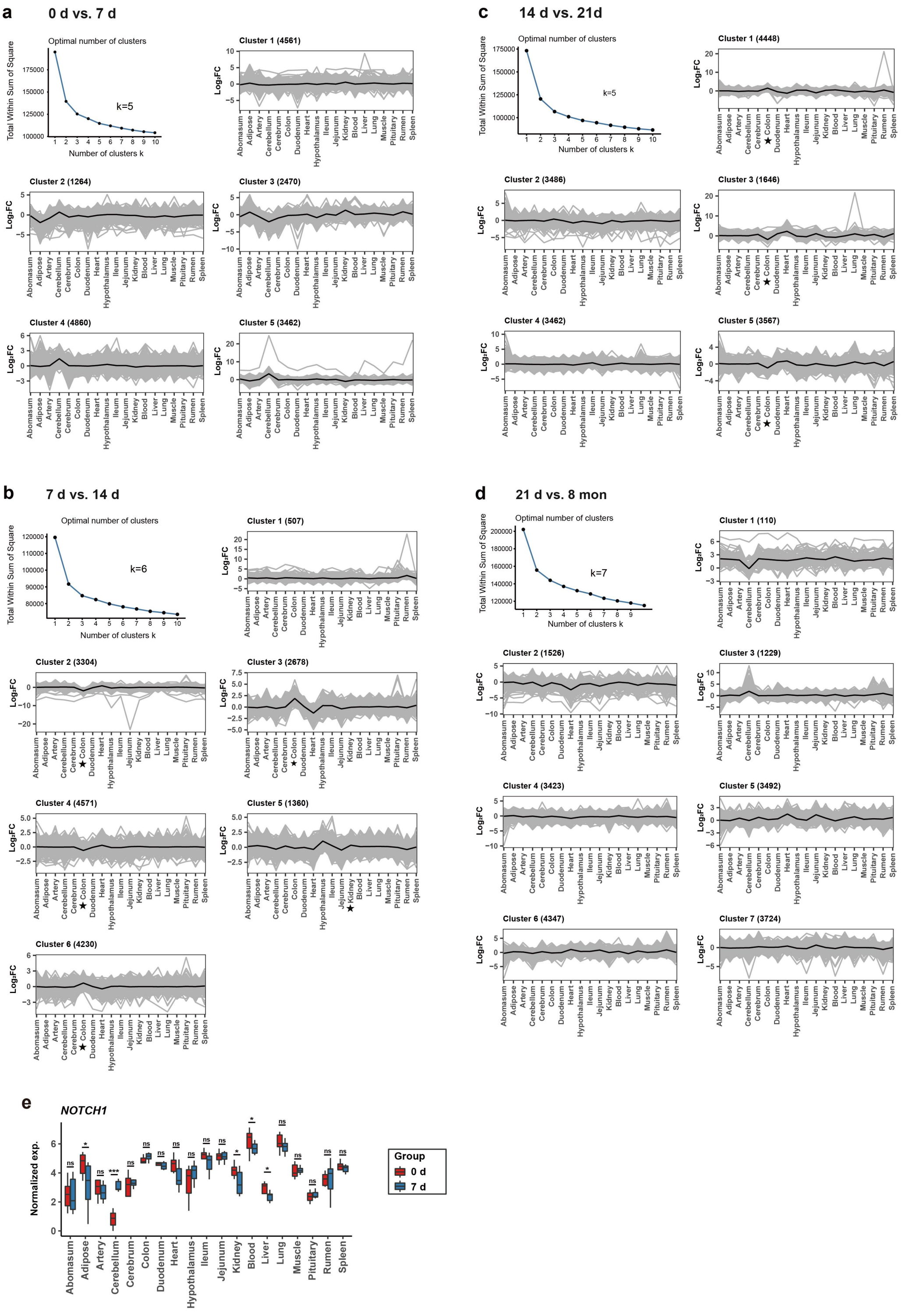


**Extended Data Fig. 8 Multi-tissue interactions in different adjacent time point comparisons.** a-d, Multi-tissue interactions in "0 d vs. 7 d" (a), "7 d vs. 14 d" (b), "14 d vs. 21 d" (c) and "21 d vs. 8 mon" (d) comparisons. Active tissues are marked with asterisks. e, The expression level of *NOTCH1* in 0 d and 7 d across tissues. The average values of $\log_2$FC for each cluster are denoted with black lines. *P* values from

Wilcoxon rank sum test, * $P < 0.05$, ** $P < 0.01$, *** $P < 0.001$.

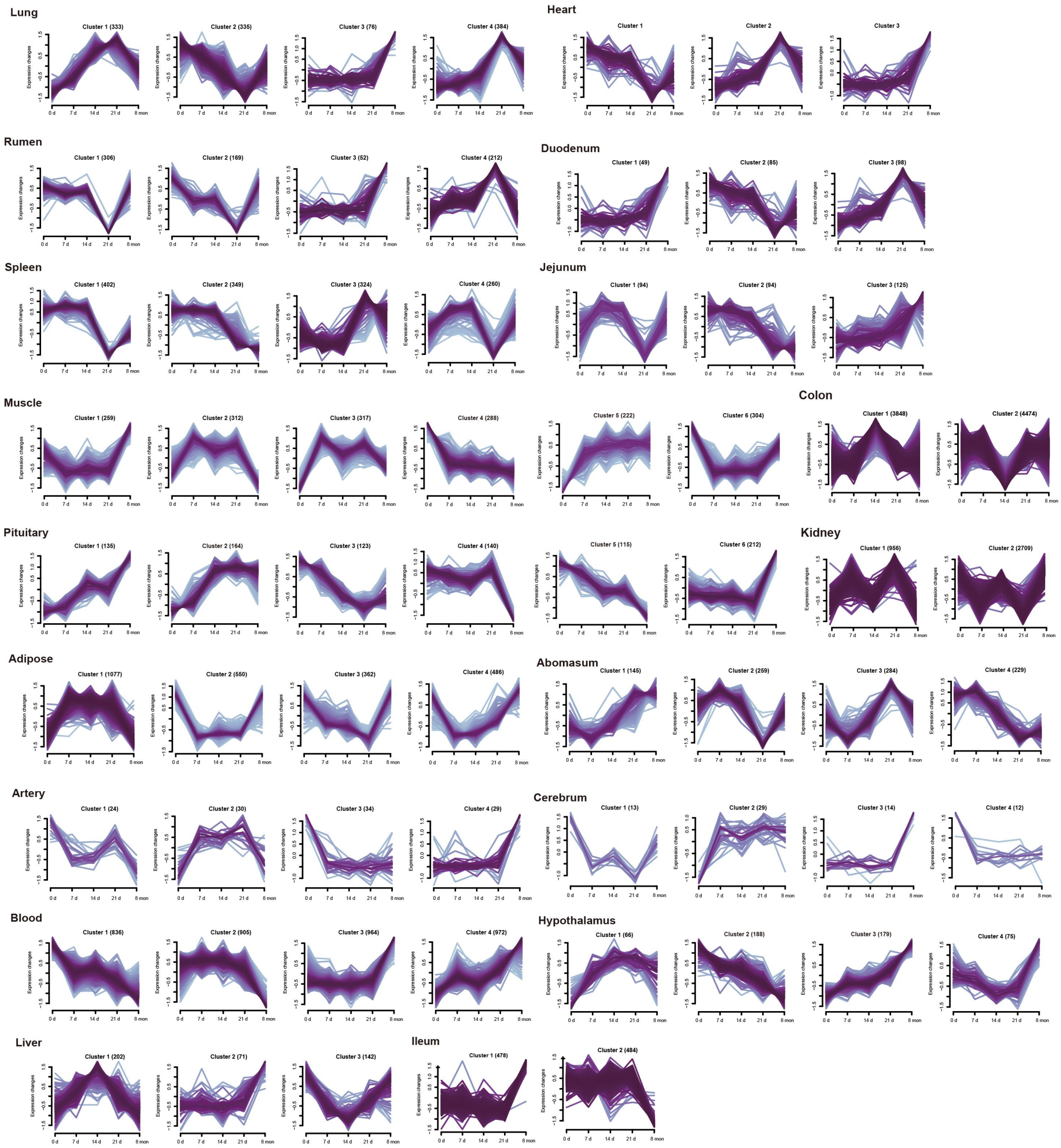


**Extended Data Fig. 9 Fuzzy c-means clustering identified gene expression patterns of DCGs across tissues.**

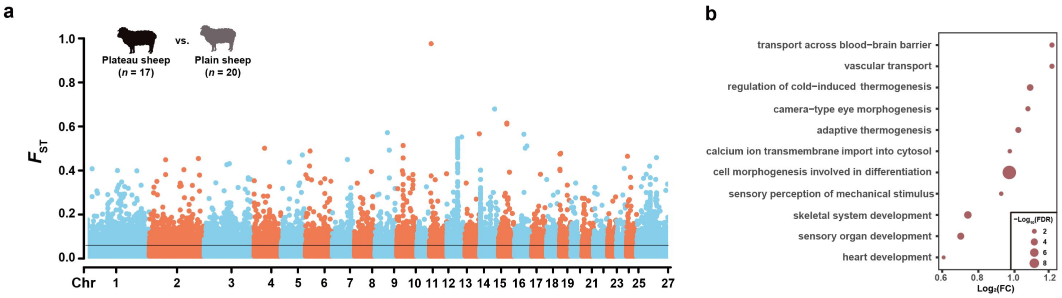


**Extended Data Fig. 10 Selective sweep analysis.** a, Genome-wide distribution of pairwise $F_{ST}$ values between plain ($n$ = 20) and plateau sheep ($n$ = 17). The top 5% significant threshold values of pairwise $F_{ST}$ ($F_{ST}$ = 0.0559) is denoted by black line. Chr, chromosome. b, GO term enrichments of candidate selective genes identified from (a). FDR < 0.05 is set as threshold.

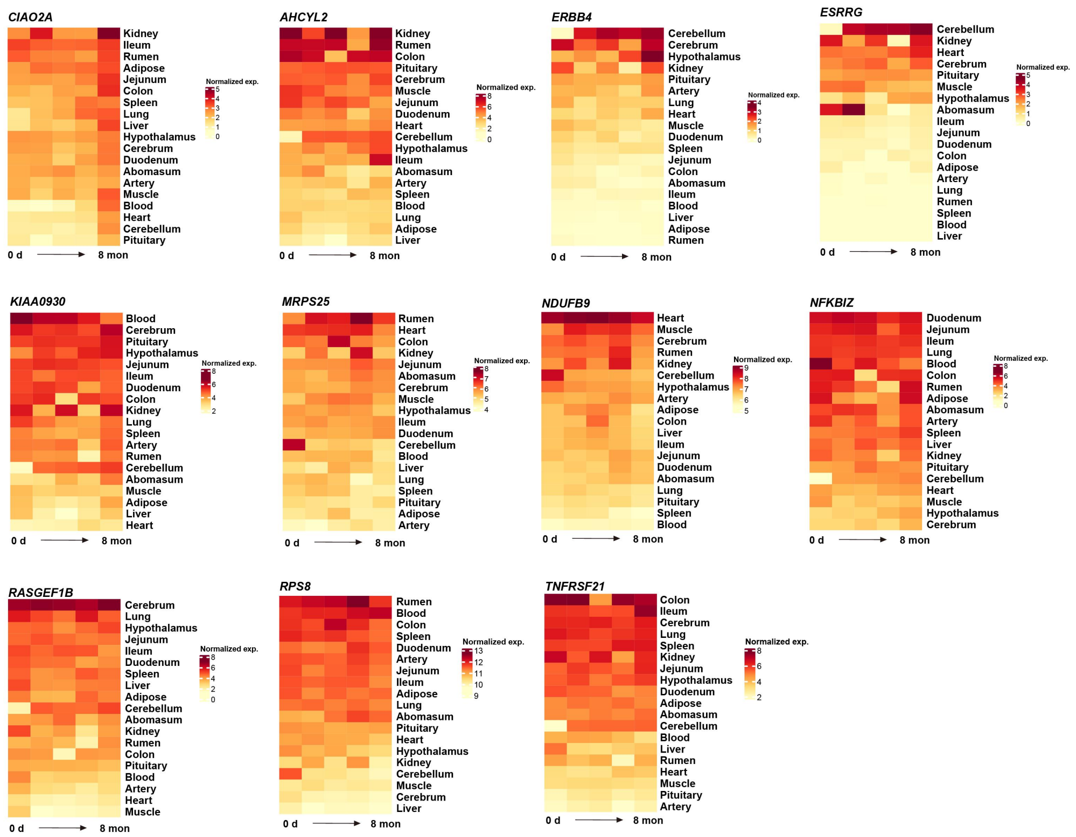


**Extended Data Fig. 11 Expression of common tissue-shared $F_{ST}$ genes.** The expression changes of common tissue-shared $F_{ST}$ genes with time across tissues.

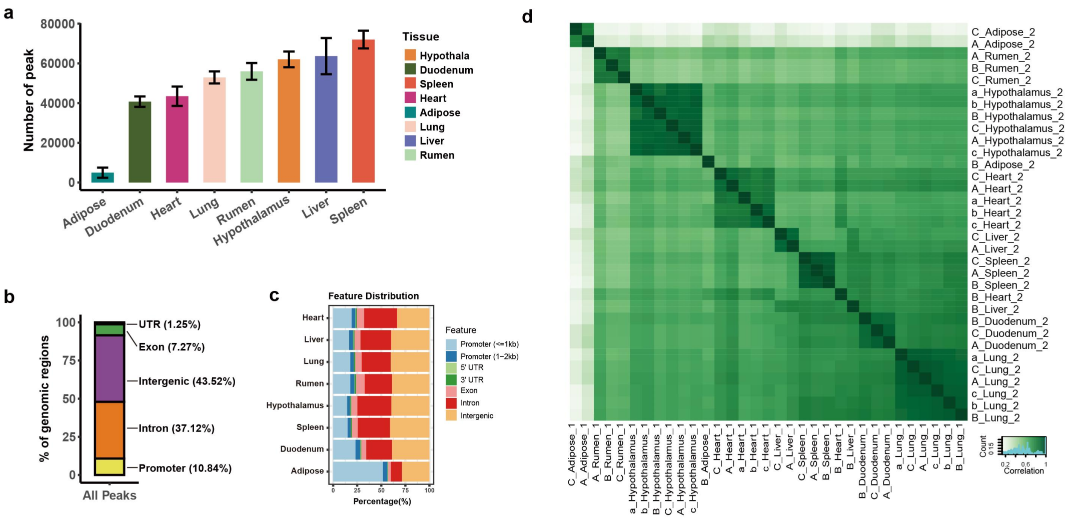


**Extended Data Fig. 12 Data summary of ATAC-Seq.** a, The number of peaks across tissues. b, The overall distribution of peaks in genomic regions. c, The distribution of peaks in genomic regions across tissues. d, Pearson's correlation between all ATAC-Seq samples based on average peak density.

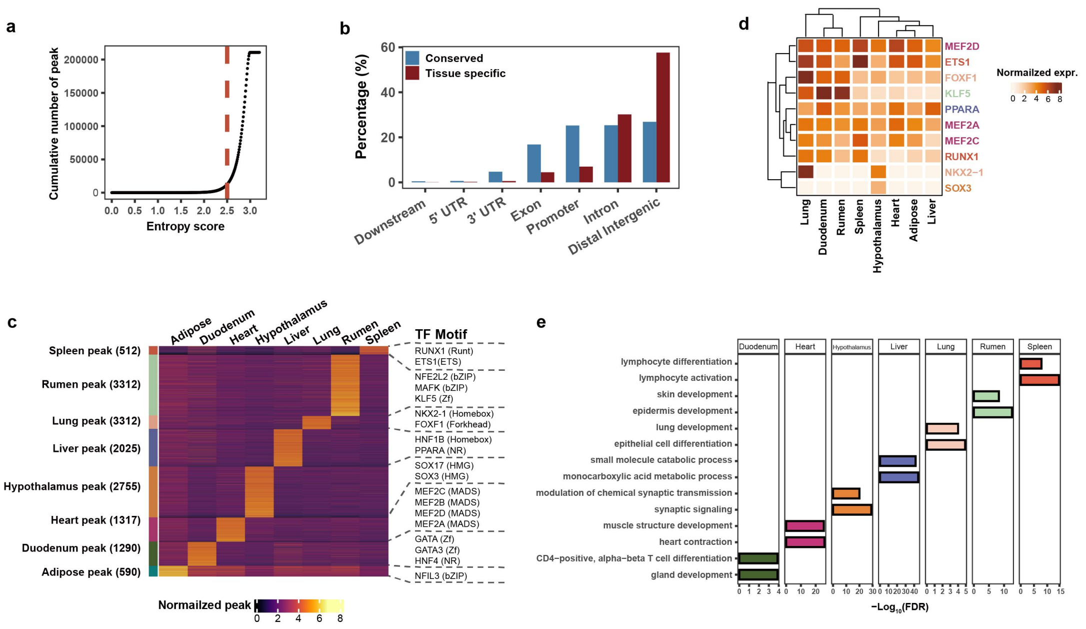


**Extended Data Fig. 13 Tissue-specific peaks and motifs.** a, The identification of tissue-specific peak based on Shannon entropy score. Entropy score = 2.5 is set as threshold and marked with red line. b, The distribution of conserved and tissue-specific peaks in genomic regions. c, Heatmap shows the signal density of tissue-specific peaks, along with the representative transcription factor (TF) motif for each tissue. *P* values come from the hypergeometric test. d, Heatmap shows expression of TF target gene across tissue. The color code for tissues is the same as in (c). e, GO term enrichments of tissue-specific peak linked genes across tissue. Two GO terms are displayed for each tissue.

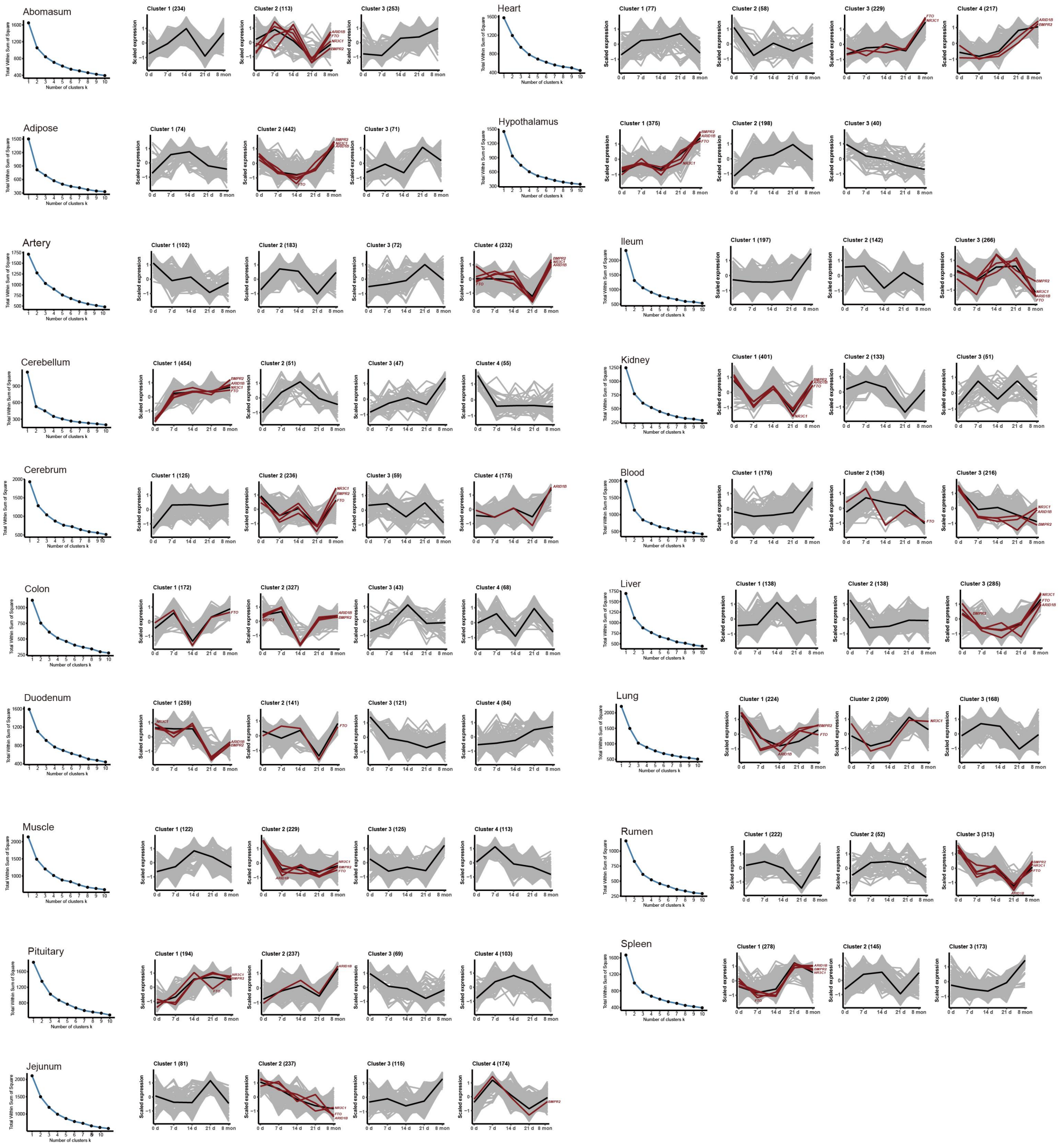


**Extended Data Fig. 14 Time-series expression for mountain sickness candidate genes of human across tissues.** We collected candidate genes ($n$ = 613) associated with mountain sickness of human. The key candidate genes of pulmonary hypertension (*BMPR2*), polycythemia (*ARID1B*), pulmonary edema (*NR3C1*) and heart failure (*FTO*) are marked with red lines. The average value of expression for each cluster are denoted with black lines.

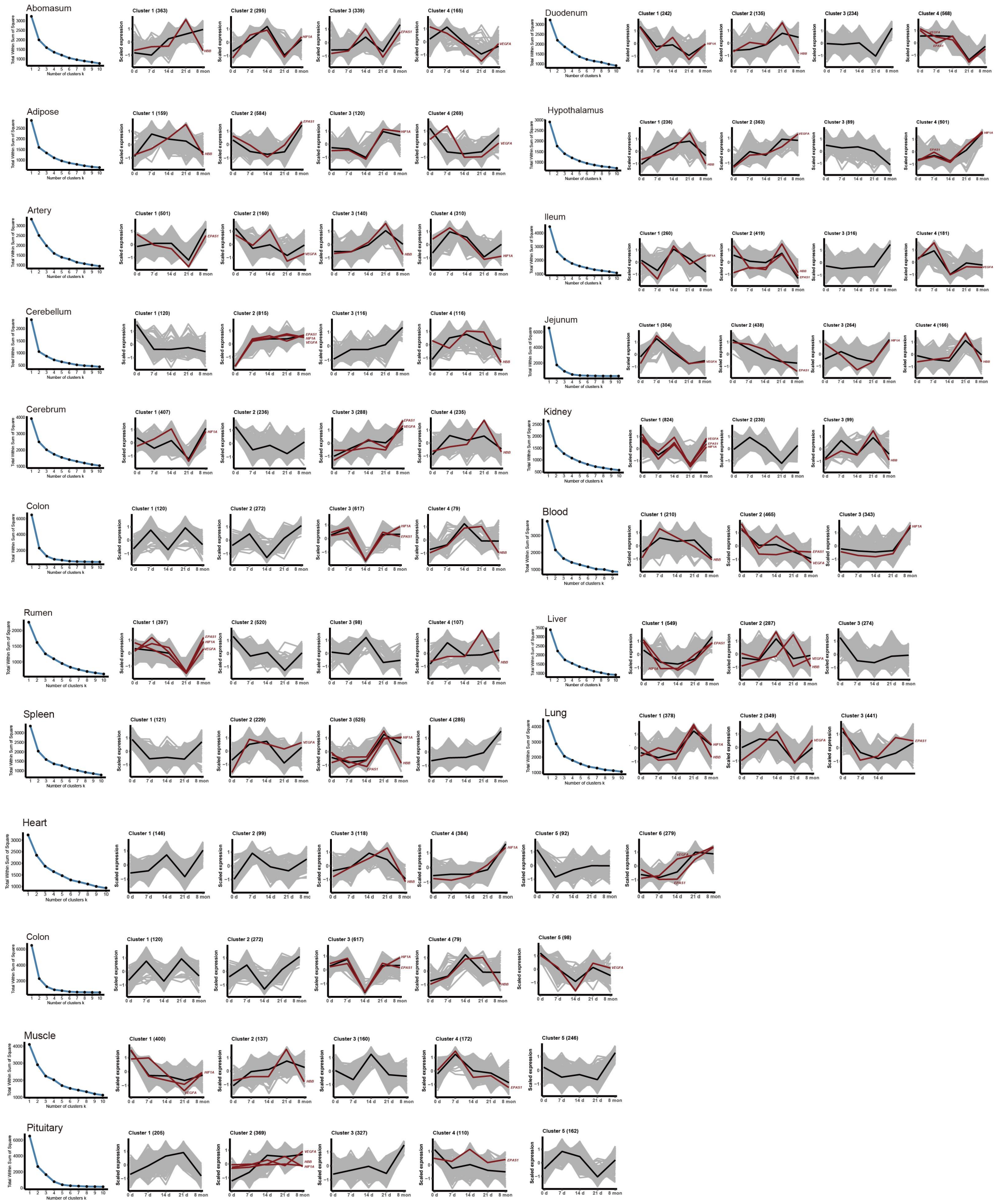

**Extended Data Fig. 15 Time-series expression for high-altitude adaptation candidate genes of human across tissues.** We collected candidate genes ($n$ =1,207) associated with high-altitude adaptation of human. The key high-altitude adaptation candidate genes *HIF1A*, *EPAS1, VEGFA* and *HBB* are marked with red lines. The average value of expression for each cluster are denoted with black lines.
